# Cadences of the Collective: Conspecific Stimulation Patterns Interact with Endogenous Rhythms to Cue Socially Mediated Response Shifts

**DOI:** 10.1101/2025.06.03.657715

**Authors:** Luke C. Larter, Colby W. Cushing, Michael J. Ryan

## Abstract

Many animals form behavioral collectives, and optimal interaction patterns often differ across social contexts. Sensory scenes generated by many interacting conspecifics are complex. Thus, maintaining socially-calibrated interaction patterns necessitates that individuals distill key features from conspecific scenes to guide continued adjustments to social fluctuations. Túngara frogs produce mating calls in choruses varying in size, and interaction patterns differ across social environments; rivals alternate their calls in smaller choruses, but increasingly overlap one another’s calls in a stereotyped fashion as choruses increase in size. We used automated playback to investigate the cues guiding this socially-mediated shift in response modes. We played conspecific stimulus calls to males at various delays relative to their own calls, preceded by various acoustic motifs mimicking conspecific interaction patterns observed across varied chorusing environments. Males almost never overlapped isolated stimulus calls at any delays. However, their probabilities of overlapping stimulus calls increased markedly when stimulus calls were preceded by motifs exhibiting intense conspecific stimulation patterns characteristic of larger choruses, especially when these stimulus calls were also presented at later delays. Thus, a multifaceted cue to social context primes varied interaction patterns on a call-by-call basis: that, in larger choruses, males experience intense conspecific stimulation during their inter-call periods, and that this stimulation extends throughout the latter reaches of their call cycles. Our results highlight that inactive phases within behavioral rhythms provide critical assessment windows for fine-tuning upcoming responses, and that behavioral rhythms act as crucial temporal filters for mapping conspecific stimulation patterns to behavioral outputs.

## Introduction

Many fitness-enhancing behaviors require successful interactions with conspecifics (Alexander, 1974). This selects for sensory tuning and cognitive abilities that allow individuals to glean salient information from conspecific cues and signals to guide adaptive responses. Conspecific signals, and potential cues, are typically complex and vary along many axes. However, not all features may be equally valuable for guiding appropriate responses, or properties of sensory or cognitive systems may constrain receivers’ abilities to use potentially informative variation along all axes. Consequently, sensory systems and social cognitive processes simplify complex incoming streams of sensory information by prioritizing processing of only the most salient features of conspecific cues and signals (Capranica, 1978; Hein, 2022; Wilczynski & Ryan, 2010). For instance, schooling and swarming behaviors can be elicited from fish and locusts by simple moving shapes (Bleichman et al., 2023; Harpaz et al., 2021). Similarly, frogs respond to highly-simplified synthetic versions of conspecific calls that retain only a few crucial features, such as species-specific note rates and broad frequency patterns (Feng et al., 1990; Gerhardt, 1974; Wilczynski et al., 1995). That coarse approximations elicit appropriate social responses demonstrates that animals’ sensory systems distill the most salient patterns from conspecific signals and cues, then rely on these essential features for guiding adaptive responses (Tinbergen, 1951).

In addition to responding appropriately to conspecifics during one-on-one interactions, animals must effectively navigate diverse social environments consisting of varied numbers and identities of conspecifics. Optimal interaction strategies often differ by social context, which selects for socially-mediated flexibility in response patterns. For instance, courting signalers increase signaling effort in more competitive signaling environments (Greenfield, 2015; Wells, 2007), combatants alter aggressive tactics based on audience composition (Matos & Schlupp, 2005), and collectively locomoting individuals alter nearest-neighbor distances in groups of different sizes (Harpaz et al., 2024; Ward et al., 2017). Sensory scenes generated in crowded social environments are invariably more complex than those generated by individual conspecifics (Bee & Micheyl, 2008; Hein, 2022; Williams et al., 2023). Thus, these examples demonstrate that, in addition to distilling salient features from individual cues and signals, animals’ sensory and cognitive systems can accomplish the even more challenging task of distilling informative features from broader conspecific scenes to guide appropriate responses across social contexts.

Scenes composed of many interacting individuals are inherently complex, as they contain features of the individuals within them, as well as emergent features and patterns arising due to interactions among these individuals. Therefore, establishing precisely which aspects of dynamic and multifaceted scenes are salient for guiding behavior can be challenging (Harpaz et al., 2024; Williams et al., 2023). Identification of salient features is complicated further by the fact that endogenous behavioral rhythms can influence how individuals interpret and respond to conspecific stimulation. This means that social cues can include a temporal component; it may not simply be the properties of conspecific stimulation that matter, but also *when* this stimulation is perceived relative to endogenous rhythms. For instance, competitive signaling interactions in *Mecapoda* katydids involve callers delaying or advancing their upcoming call in response to a rival’s call, and these temporal adjustments differ depending on when this rival’s call is perceived relative to endogenous calling rhythms (Hartbauer et al., 2005; Sismondo, 1990). Similarly, juvenile zebrafish swimming dyadically alter the intervals between successive locomotion bursts as a function of the delay between their most recent burst and their partner’s response burst, generating temporally-coupled bursting dynamics (Amichay et al., 2024). Beyond one-on-one interactions, behavioral rhythms can also influence individuals’ responses to emergent regularities in conspecific scenes within behavioral collectives. For instance, marching locusts and schooling zebrafish alternate between phases of rapid and slowed locomotion, and are more visually responsive to the predominant movement directions and velocities of group-members during slowed phases (Aidan et al., 2024; Ariel et al., 2014; Harpaz et al., 2017). Thus, in many collective contexts, salient cues guiding socially mediated flexibility are complex and multi-dimensional, with responses being guided by how emergent conspecific stimulation patterns interact with, and are filtered through, the temporal prism of individuals’ endogenous responsiveness rhythms (Larter & Ryan, 2025).

Acoustic chorusing is a collective behavioral context in which the rhythmicity of behavior, and socially-dependent flexibility, have been extensively studied; many playback experiments have probed the temporal coupling functions facilitating dyadic calling interactions (Greenfield, 1994a; Klump & Gerhardt, 1992; Larter and Ryan, 2025), and many have investigated how callers alter calling strategies across varied social environments (Greenfield, 2015; Wells, 2007). However, no studies have experimentally investigated the role that interactions between emergent regularities in conspecific scenes and individuals’ endogenous call rhythms play in guiding flexible signaling strategies. Here, we utilized an automated playback paradigm to investigate whether such an interaction is responsible for cueing socially mediated shifts in response modes in túngara frogs. This species shows two largely discrete response types; a male can alternate with a rival’s call without overlapping it, or can overlap it in a stereotyped way (Larter & Ryan, 2024a; described in the Methods). Prevalences of these different response types strongly correlate with chorus size, with alternation predominating in smaller choruses and overlap becoming increasingly prevalent as choruses grow beyond 3 males. Furthermore, observations hint that overlap is driven by an interaction between endogenous calling rhythms and broader conspecific stimulation patterns (Larter & Ryan, 2024c).

To explore this experimentally, we played conspecific stimulus calls to males at various delays relative to their own calls, and preceded these stimulus calls with a variety of acoustic motifs that mimicked conspecific stimulation patterns typical of different social environments. We hypothesized that the probability that a male overlapped the stimulus call would be influenced by an interaction between the motif preceding the stimulus call and the delay at which he encountered it relative to his most recent call. We predicted that: ***i)*** stimulus calls preceded by motifs typical of more crowded choruses would have higher probabilities of being overlapped, ***ii)*** stimulus calls encountered at later delays throughout males’ call cycles would have higher probabilities of being overlapped (Larter & Ryan, 2024c), and ***iii)*** that these factors would interact, such that overlap probability would be increased synergistically when stimulus calls were both preceded by crowded chorus motifs and encountered at later delays. Furthermore, males exhibit socially-dependent flexibility in call elaboration patterns (Bernal et al., 2007; Larter & Ryan, 2024a), and response latencies to offsets of rivals’ calls when alternating (Larter & Ryan, 2024a), and we predicted that analogous synergistic effects would also underpin flexibility in these aspects of calling behavior.

## Materials and Methods

### Túngara frogs (*Physalaemus* (=*Engystomops*) *pustulosus*)

Túngara frog males form choruses in shallow puddles and pools. Choruses can be dense, and density can vary within the same area on the same night (Bernal et al., 2007). This species’ calls begin with a ‘whine’, a descending frequency sweep, to which can be added 1 or more broadband ‘chuck’ notes (Figure 1; Ryan, 1985). Calls including chucks (‘complex calls’) are 5-fold more attractive to females than whines alone (‘simple calls’) (Ryan et al., 2019). Median call periods (time elapsing between the onset of one call and the next for a given caller) are ∼1.7s for chorusing males (Larter & Ryan, 2024a).

**Figure 1:**
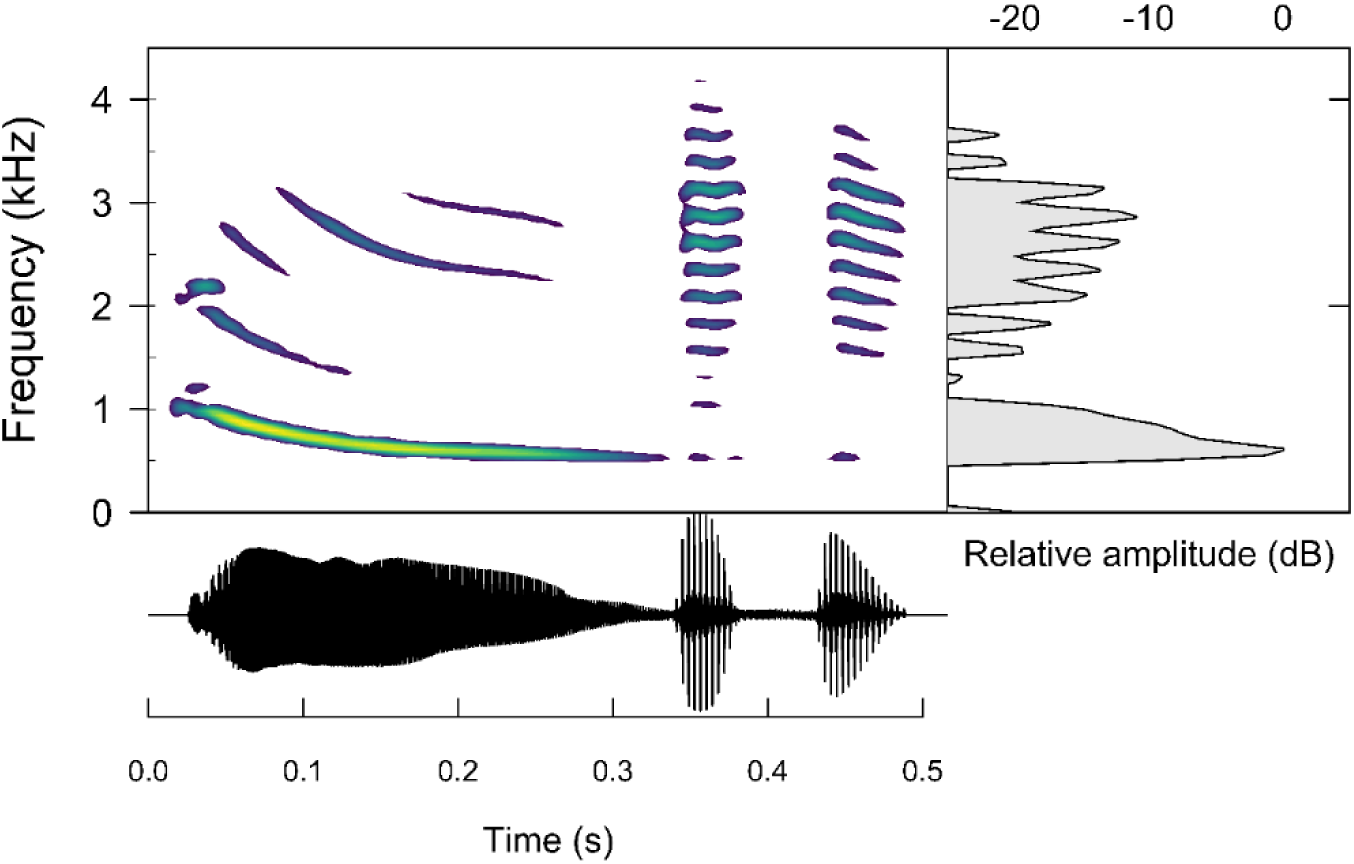
A spectrogram (upper), waveform (lower), and power spectrum (right) of a complex túngara frog call (‘seewave’ R package, window length 1024, overlap 90%). Brighter colors denote higher intensities. This call begins with a continuous ‘whine’ note and ends with 2 ‘chuck’ notes.

Calling interactions among rivals are mediated by a ‘gap-detection’ strategy. Here, calls are triggered when males experience an abrupt reduction in inhibition by acoustic stimulation, as this is indicative of the onset of a low-interference ‘gap’ in ongoing chorus noise into which an impending call can be inserted (Larter & Ryan, 2024b; Larter and Ryan, 2025). However, interaction patterns differ by chorus size. In choruses of 3 or fewer males, rivals alternate their calls without overlapping. However, above this threshold, call overlap among rivals becomes increasingly prevalent. This overlap is highly stereotyped, with whine onsets of the overlapping following calls beginning just before the chucks of the overlapped leading calls (Larter & Ryan, 2024a). For following males, overlap of this form induces lower attractiveness costs than any other potential form of overlap while also reducing the attractiveness of the overlapped leading call, making this a beneficial interaction pattern when overlap becomes unavoidable in larger choruses (Larter & Ryan, 2024a).

Furthermore, alternation and stereotyped call overlap seem to be facilitated by the same gap-detection mechanism. Amplitude and frequency decrease concurrently throughout whines, and male sensory systems are tuned such that this concurrent decrease generates a moderately steep release from inhibition shortly after whine onsets which can trigger calls under certain circumstances (Larter & Ryan, 2024b; illustrated in Larter and Ryan 2025). Thus, alternation in smaller choruses occurs because the stark release from inhibition accompanying the offsets of rivals’ calls to silence is a highly salient call-trigger. Conversely, stereotyped overlap in larger choruses occurs as the moderately steep release from inhibition occurring just after whine onsets becomes an increasingly salient call-trigger as choruses grow larger (Larter & Ryan, 2024a, 2024b). Thus, the salience of this latter call-trigger varies across social environments, which drives these varied interaction patterns. Furthermore, chorusing males were more likely to overlap rivals’ calls encountered later throughout their call cycles, suggesting salience of this latter call-trigger is also mediated by males’ endogenous calling rhythms (Larter & Ryan, 2024c).

Other aspects of male calling behavior also correlate with chorus size. In larger choruses, males increase call elaboration by appending more chucks to calls (Bernal et al., 2007; Larter & Ryan, 2024a), and decrease response latencies to rivals’ call offsets when alternating with them (Larter & Ryan, 2024a).

### Playback Experiments

#### Experimental Subjects

During July and August 2024, we collected males as members of amplectant pairs from urban breeding sites around Gamboa, Panama, near the Smithsonian Tropical Research Institute grounds. We separated males from their associated females for trials and, following trials, weighed males (g), and gave them a unique toe clip to avoid retesting. We then reunited males with their females and returned pairs to their capture locations later that same night. Morphological and environmental data for 3 of 39 males were lost, so we assigned these males median weights and water temperatures.

All research was permitted by the Government of Panama (ARB-130-2023), approved by STRI-ACUC (SI-24020-1) and UT Austin IACUC (AUP-2022-00012), and followed the Guidelines for Use of Live Amphibians and Reptiles in Field and Laboratory Research (Beaupre et al., 2004).

#### Stimuli

For stimulus calls, we chose three calls from previous recordings, each made by a different male as they called together as members of the same 6-male chorus (Larter & Ryan, 2024c). These males differed widely in their probabilities of being overlapped by their rivals, as ascertained by extracting males’ random intercepts from the generalized linear mixed-effects model (GLMM) predicting overlap probability in that paper. Prior to doing so, we removed all pertinent fixed effects to concentrate all inter-male variation in this random intercept. ‘Stimulus call IDs’ denote these relative differences in previously-observed overlap probabilities (OP) between calls: ‘low OP’ (P(overlapped) = 0.18), ‘intermediate OP’ (P(overlapped) = 0.52), and ‘high OP’ (P(overlapped) = 0.84). Durations of these calls correlated with overlap probabilities (low OP = 0.428s < intermediate OP = 0.484s < high OP = 0.519s). However, looking more closely at overlap patterns revealed that differences in overlap probability were not driven by differences in duration per se, which we discuss in the Results. We also chose a fourth call, by a male from a different chorus who also had a low probability of being overlapped by his rivals (P(overlapped) = 0.12). We refer to this as the ‘interim call’, and responses to this call were not tested. Rather, the interim call was played to males at a fixed delay in between stimulus call presentations, to provide a standardized non-overlapping interaction before every stimulus call presentation, and we used interim calls to construct acoustic motifs preceding stimulus calls (Figure 2).

**Figure 2:**
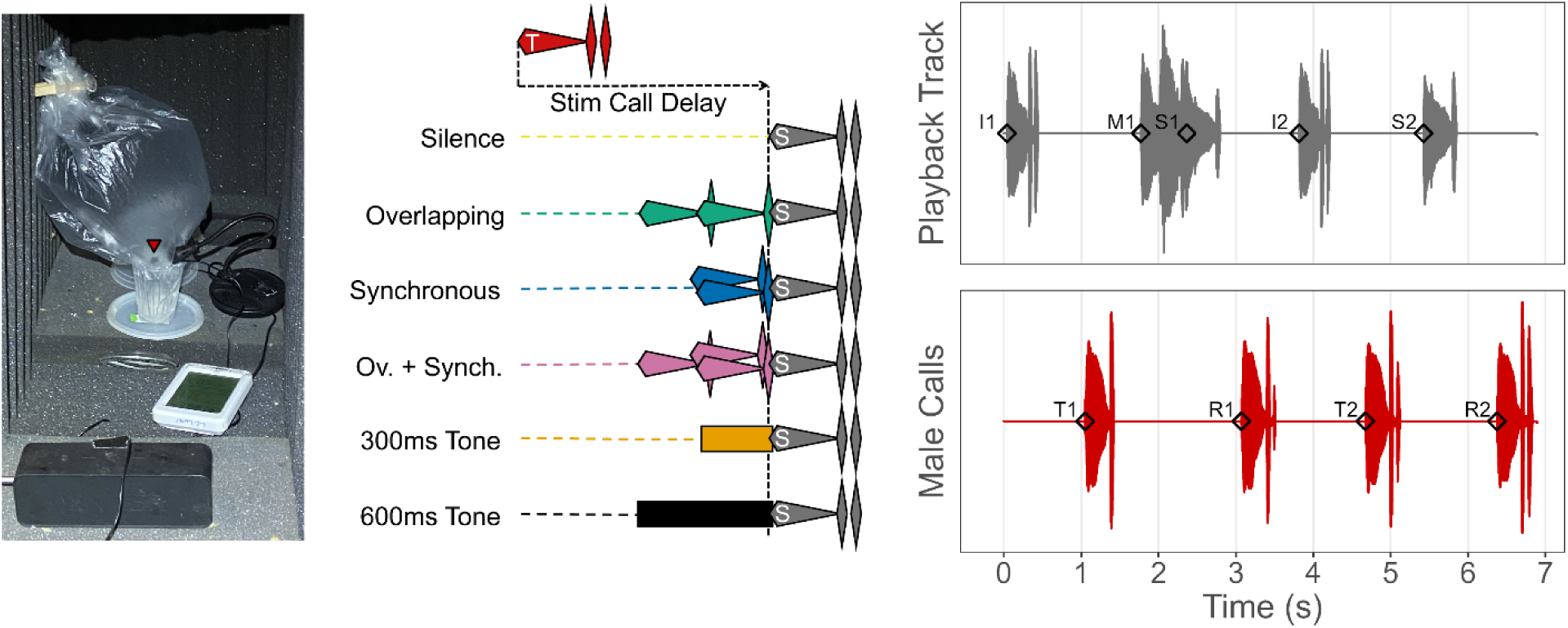
***Left***: A male (red arrow) in our automated playback setup. The microphone registered his call onsets which triggered stimuli to be played from the speaker in the foreground. ***Middle***: Illustrations of acoustic motifs that preceded stimulus calls. ‘T’ call represents the male’s trigger call, and ‘S’ calls represent stimulus calls that could be played back at an array of stimulus delays, and could be preceded by any of these motifs. ***Right***: An example of a male (*lower trace*) interacting with automated playback (*upper trace*). In the playback track, I1 and I2 denote onsets of ‘interim calls’, S1 and S2 denote onsets of ‘stimulus calls’, and M1 the onset of the ‘motif’ preceding S1 (here, overlapping+synchronous). S2 here is preceded by silence. In the male response track, T1 and T2 denote onsets of ‘trigger calls’ that initiate stimulus call playback, and R1 and R2 denote onsets of ‘response calls’ to these stimulus calls. Key variables of interest are visible here: **i)** ‘**Stimulus delay**’: the time elapsing between the onset of a <u>trigger call</u> and the onset of the consequent <u>stimulus call</u> (e.g., S1 - T1 (*not* M1 - T1)); our temporal predictor variable of interest. **ii)** ‘**Response delay**’: time elapsing between the onset of a <u>stimulus call</u> and the onset of the subsequent male <u>response call</u> (e.g., R1 - S1); a temporal response variable of interest. **iii)** ‘**Response period**’: time elapsing between the onset of a male’s <u>trigger call</u> and the onset of his subsequent <u>response call</u> following stimulus presentation (e.g., R1 - T1); another temporal response variable of interest, equivalent to ‘stimulus delay’ + ‘response delay’. **iv)** ‘**Trigger call elaboration**’: the summed amplitude, relative to the peak amplitude of the associated whine, of all chuck notes appended to the trigger call that initiated playback of the stimulus call to which the male was currently responding. Increasing call elaboration is visible throughout the lower trace.

We presented stimulus calls to males at various delays, either in isolation (preceded by silence) or preceded by acoustic motifs composed of combinations of overlapping calls, synchronous calls, or tones (Figure 2). Overlapping and synchronous call motifs mimicked analogous interactions observed in larger túngara frog choruses (Larter & Ryan, 2024a, 2024c). Motifs were constructed using our interim call with its second chuck removed. This call previously had a low probability of being overlapped, suggesting males would be stimulated by our motifs but would seldom overlap them, thereby focusing their responses on the stimulus calls of interest that followed motifs. Additionally, to disentangle the effects of more specific features of conspecific stimulation vs. generic inhibition by intense acoustic stimulation, we included motifs consisting of 1000Hz tones, which strongly inhibit calling (Larter & Ryan, 2024b). Thus, stimulus calls could be preceded by: ***i***) silence (i.e., isolated stimulus calls), ***ii***) two overlapping calls (610ms in duration), ***iii***) 2 synchronous calls (370ms), ***iv***) a combination of overlapping and synchronous calls (denoted ‘overlapping+synchronous’ to avoid confusion; 610ms), ***v***) a 610ms 1000Hz tone, or ***vi***) a 310ms 1000Hz tone (Fig 1). The last 10ms of motifs overlapped with the first 10ms of stimulus calls, to generate smoother transitions free from low-amplitude dips, with motifs extending up to 600ms before stimulus call onsets.

Stimulus calls preceded by silence were played at 8 ‘stimulus delays’ relative to the male’s trigger call onset: 0.45s (the duration of a typical call: (Ryan & Rand, 2003)), 0.6s, 0.75s, 0.9s, 1.05s, 1.2s, 1.35s, and 1.5s. However, males sometimes called spontaneously just prior to onsets of stimulus calls presented at later delays, which also gave us responses to stimulus calls that overlapped trigger calls (though these responses were biased towards males with the fastest intrinsic call rates; discussed in the Results). Conversely, as acoustic motifs extended up to 0.6s before stimulus call onsets, we could only play stimulus calls preceded by them at later stimulus delays: 1.05s, 1.2s, 1.35s, and 1.5s. To clarify, stimulus delays denote the delay between the onset of the subject’s trigger call and the onset of the consequent stimulus call, not the onset of the motif preceding it (see Figure 2). All stimulus call/stimulus delay/motif combinations were played to e ach male in a unique randomized order within a single trial, with each combination appearing 5 times (490 total stimulus presentations in a full trial).

#### Automated Playback Trials

Trials took place in a darkened room. During trials, a male called while floating on the surface of water contained in a 6.5cm-diameter cup within an acoustically-transparent enclosure (Figure 2). Water temperature ranged from 26.4 - 28.6C. A JOUNIVO USB microphone registered the male’s call onsets to facilitate interactions with our automated playback program. Playback stimuli were played from an ANKER SoundCore speaker positioned 38cm from the center of the male’s cup. We calibrated this speaker such that peak amplitudes of whines of stimulus and interim calls, and 1000Hz tones, registered at 82dB SPL (re. 20 µPa) (the amplitude of a typical call from ∼1.3m away) at the center of the cup. This setup was surrounded by acoustically-insulating foam.

The microphone and speaker were connected to a laptop running our automated playback program via a Jupyter (v7.0.8) notebook coded in Python (v3.11.7), that utilized functionalities from the PyAudio package (v0.2.14). This allowed us to play stimulus calls to males at our desired array of stimulus delays. A brief outline of playback trials is as follows:

1) At the start of trials, males were stimulated to call via 1 minute of playback of interim calls.
2) The onset of the male’s call in response to the final interim call in this sequence then triggered playback of the first stimulus call in the trial.
3) This stimulus call was played to the male at the desired stimulus delay, and the onset of his call in response to this stimulus call then triggered an interim call to be played at a fixed delay of 0.75s.
4) The onset of the male’s call in response to this interim call then triggered the next stimulus call in the sequence. Then, back to 3.

Steps 3 and 4 then alternated, with stimulus and interim calls presented alternately for remaining stimuli (Figure 2). This ensured a ‘palette-cleansing’ alternation interaction with an interim call between each stimulus call presentation, to reduce carry-over effects. Full trials lasted ∼35 minutes, though we ended trials early if males ceased calling for several minutes.

Timestamps recorded by our automated playback program for trigger, response, and stimulus call onsets were not sufficiently precise or consistent for analysis (+-50ms). Thus, we also recorded the subject and playback speaker via separate Synco LavS6R tie-clip microphones onto a Zoom F6 recorder. We used precise timestamps extracted from these recordings using functions from the ‘Librosa’ Python package (McFee et al., 2015) for analysis.

## Data Analysis

### Response Delay and Response Period Curves

Phase response curves are commonly used to investigate temporal responses of calling males to rivals’ calls encountered at different points throughout their call cycles (Greenfield, 1994a; Larter and Ryan, 2025). We constructed analogous curves to characterize male temporal responses to stimulus calls across our range of stimulus delays. However, rather than normalizing stimulus and response delays to phase angles, we modeled them in absolute time and accounted for inter-male variation using random effects.

We modeled temporal responses to stimulus calls presented at different stimulus delays (***Response_Delay_GAMM***: response delay ∼ stimulus delay), and the consequences of these responses for male calling rhythms (***Response_Period_GAMM***: response period ∼ stimulus delay) using generalized additive mixed-effects models (GAMM) with inverse-gaussian distributions (‘mgcv’ R package: Wood, 2001). Here, we only included responses to stimulus calls preceded by silence, and excluded responses where males overlapped stimulus calls rather than alternating with them (only 150/4399 (3.5%) of responses). This yielded n = 4,249. For *Response_Delay_GAMM*, we included response delay as the response variable and, for *Response_Period_GAMM*, we included response period as the response variable. Otherwise, these models were identical. We included stimulus delay as a predictor variable and modeled this relationship with a nonlinear smooth containing 10 knots. We also included a random intercept and random smooths for individual males, to capture inter-male variation. Additionally, we included linear covariates expected to influence responses; *i)* the onset time of response calls within trials (to control for potential fatigue or habituation effects), *ii)* the call elaboration score of the trigger call initiating each playback (standardized within males; to control for within-male variability in general arousal levels throughout trials), *iii)* stimulus call ID as an unordered factor (low OP, intermediate OP, and high OP), *iv)* male weight (in grams), and *v)* water temperature at trial beginning (degrees C). We included random slopes for most linear variables, except for weight and water temperature for which males only experienced a single value. Throughout our analysis, we checked model assumptions using the ‘DHARMa’ R package (Hartig, 2022); all models and checks can be rerun using the associated R code.

### Effects of Preceding Motifs on Responses to Stimulus Calls

We used mixed effects models (‘lme4’ R package: Bates et al., 2007) to investigate how male responses to stimulus calls were influenced by interactions between stimulus call ID, the motifs preceding them, and the stimulus delays at which they were played. To minimize risks of type 1 errors (falsely significant results), we did not simplify the fixed-effect structure of models (Schielzeth & Forstmeier, 2009), and we fit the maximal random-effects structures supported by the data (Barr et al., 2013). We included random intercepts for subject identity nested within testing block, and initially included correlated random slopes for all predictor variables (except mass and water temperature for which each subject only experienced a single value). However, when correlated random slopes for certain variables were not supported, we removed correlation terms, then removed these random slopes entirely if they remained unsupported. We standardized all continuous predictor variables ((x-mean(x))/S.D.(x)) prior to analysis, with trigger call elaboration score being standardized within males to account for the large degree of intermale variation in typical elaboration levels (Larter & Ryan, 2024a).

#### Overlap_GLMM

To investigate the factors responsible for inducing overlapping responses, we built a mixed effects logistic regression model with whether the response call onset occurred anywhere during the stimulus call or not as the response variable (n = 12,153). As predictor variables, we included stimulus delay, motif preceding the stimulus call, and stimulus call ID (low OP, intermediate OP, high OP). As we anticipated these factors would interact, we included all 2-way interactions between these variables. We also included the same control covariates as in our GAMMs: the onset time of male response calls, trigger call elaboration score, male weight, and water temperature.

#### Response_Delay_LMM

To investigate the factors influencing response delays, we built a log-linear mixed effects model with log(response delay) as the response variable. Here, we only included the subset of responses that did not overlap stimulus calls, i.e., alternated with them, allowing us to investigate response delays relative to an identifiable call-trigger (stimulus call offsets) (n = 8,863). We included the same predictor variables and interactions as in *Overlap_GLMM*.

#### Within_Male_Call_Elaboration_LMM

Typical call elaboration levels vary among males (Larter & Ryan, 2024a), and males adjust call elaboration levels in response to chorus density (Bernal et al., 2007). In contrast to temporal responses which often vary greatly from call-to-call, call elaboration levels remain relatively stable over long stretches of calling (Bernal et al., 2009). To investigate how our factors of interest influenced short-term within-male changes in call elaboration, we built a linear mixed effects model with response call elaboration score as the response variable (n = 12,153) and trigger call elaboration score as a control covariate. Importantly, both trigger and response call elaboration scores were standardized using their collective mean and standard deviation. Thus, this model reveals to what degree different playback stimuli combinations induced an increase or decrease in response call elaboration scores relative to the trigger call immediately prior to them, adjusted to account for differences in males’ typical call elaboration levels throughout trials. We included the same predictor variables and interactions as in *Overlap_GLMM* except that, as we were modeling within-male call elaboration patterns, we did not include weight and water temperature since each male only experienced a single value and their effects would be subsumed by our procedure of standardizing response call elaboration score within-males.

#### Between_Male_Call_Elaboration_LMM

To investigate the influence of phenotypic (body weight) and environmental (water temperature) factors on absolute call elaboration levels, we built a log-linear mixed effects model with log(unstandardized response call elaboration score) as the response variable. Log-linear models cannot handle zeroes, so we removed 12 calls with response call elaboration scores of 0 (n = 12,141). Here, we included the same predictor variables as *Overlap_GLMM*, except that we removed trigger call elaboration score, as we simply wanted to model mean response call elaboration scores rather than elaboration changes relative to trigger calls.

### AI Declaration

AI was used as a search tool (Microsoft Copilot, Consensus), and for generating code templates as starting points for figures (Microsoft Copilot).

## Results

Unless otherwise stated, predicted probabilities presented here for focal predictors of interest are the estimated marginal means, with non-focal continuous covariates held at mean values and predictions averaged across all levels of non-focal categorical covariates (via ‘sjPlot’ R package: Lüdecke, 2021). We used an α of 0.05 to determine statistical significance.

### Response Delay and Response Period Curves

This species’ response delay curve (***Response_Delay_GAMM***) is roughly piece-wise linear, beginning with a near-linear decrease in response delays to stimulus calls played at short stimulus delays that overlapped male trigger calls (0 to 0.45s) (but see Figure 3 caption for caveats), and flattening out for all later, non-overlapping, stimulus delays (Figure 3). Though males exhibited variation in overall response curve shape (Supplemental Information SI3), this general shape predominated, and most variation was in curve intercepts. Other variables significantly influenced response delays (Table 1). Stimulus call ID influenced response delays (predicted response delay (s) [95% CI]: low OP, 0.86 [0.81, 0.9]; intermediate OP, 0.94 [0.89, 0.99]; high OP, 0.77 [0.74, 0.81]), higher trigger call elaboration scores decreased response delays slightly (-1SD, 0.87 [0.83, 0.91]; mean, 0.85 [0.81, 0.89]; +1SD, 0.84 [0.8, 0.88]), and larger males exhibited shorter response delays (1.3g body weight, 0.91 [0.85, 98]; 1.7g, 0.85 [0.81, 0.89]; 2.1g, 0.79 [0.73, 0.85]). The effects of water temperature (26.5C, 0.86 [0.77, 96]; 27.5C, 0.85 [0.81, 0.89]; 28.5C, 0.85 [0.77, 0.93]), and call onset time within trials (5 minutes, 0.86 [0.82, 9]; 17.5 minutes, 0.85 [0.81, 0.89]; 30 minutes, 0.85 [0.8, 0.89]) were negligible and non-significant. Results for ***Response_Period_GAMM*** were equivalent, just in the slightly different context of response periods rather than response delays. The flat latter arm of the response delay curve (*Response_Delay_GAMM*) yielded a response period curve (*Response_Period_GAMM*) that increased linearly in a 1:1 manner for all stimulus delays beyond those that overlapped trigger calls (Figure 3).

**Figure 3:**
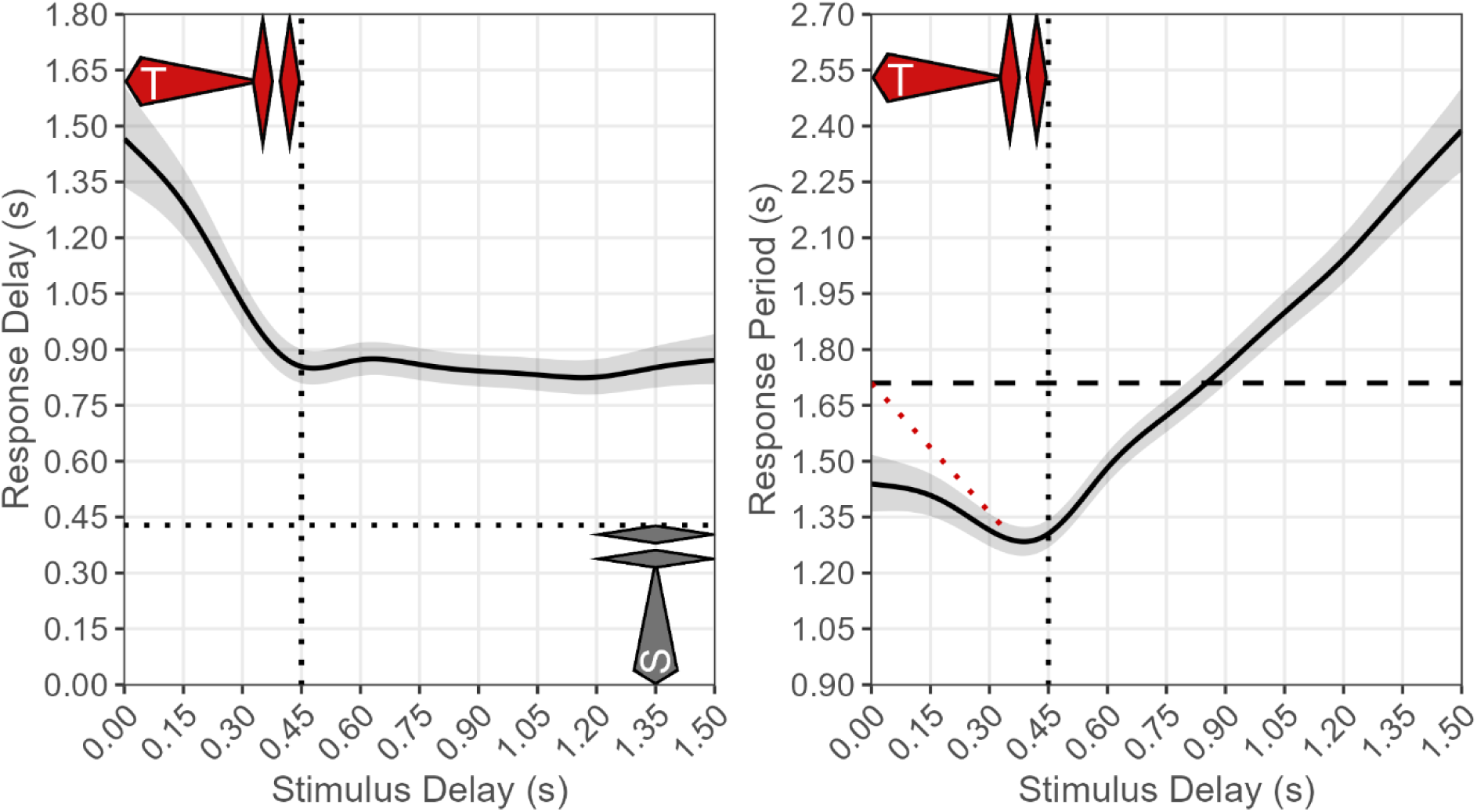
***Left***) *Response_Delay_GAMM* results: Average response delays to isolated low OP stimulus calls encountered at various stimulus delays. Red call approximates a typical trigger call (Ryan & Rand, 2003), with stimulus delays to the left of the vertical dotted line representing stimuli that likely overlapped trigger calls to various degrees. The gray call represents the stimulus call. Plotting responses to a single stimulus call ID allows us to visualize responses relative to a specific stimulus call duration (horizontal dotted line); low OP was chosen arbitrarily. ***Right***) *Response_Period_GAMM* results: Average response periods arising due to response delays shown in left figure. The horizontal dashed line at 1.71s depicts the expected call period (ECP: median call period observed during dyadic chorusing: Larter & Ryan, 2024a). Thus, when the curve runs below this line, this indicates a shortening of call periods relative to the ECP, and vice versa. Intercepts of such curves (e.g. phase response curves) are typically equivalent to the ECP (see examples in Larter and Ryan, 2025). That the intercept here is below the ECP likely arises because we did not systematically present stimulus calls at these short, overlapping, stimulus delays (<0.45s). Rather, we obtained these data opportunistically when males called spontaneously shortly before onsets of stimulus calls presented at later delays. Our maximum delay was 1.5s, meaning datapoints to the left of the vertical dotted line (in both plots) are biased towards a subset of males who commonly exhibited very short call periods <1.5s (plotted intercept is just under 1.5s) (SI4). The dotted red line intersecting the ECP is what would be expected logically, and based on similar curves for other frogs (Lemon & Struger, 1980), and so likely approximates what this first arm would look like with more systematic sampling.

**Table 1:**
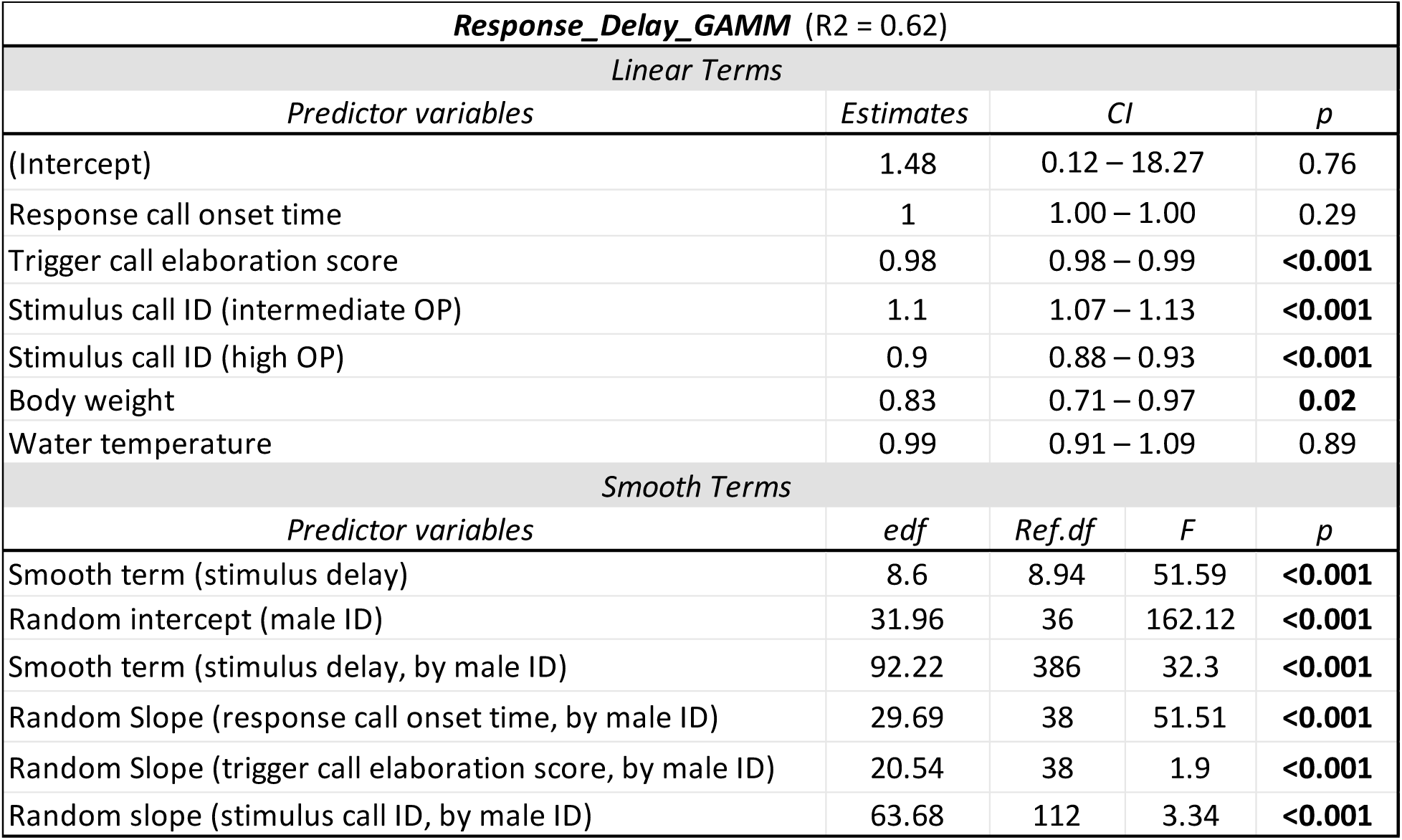
Results of *Response_Delay_GAMM* (response delay ∼ stimulus delay). Results for *Response_Period_GAMM* were similar (SI3). Predictor variables for GAMMs were not standardized prior to inclusion, with the exception of trigger call elaboration score which was standardized within males.

### Effects of Preceding Motifs on Responses to Stimulus Calls

For our mixed-effects models, model summaries are presented in Appendix 1, marginal effects of our main interactions of interest are visualized in Figure 4, and marginal effects of control variables are presented in Table 2. See SI5 for random effects summaries. p-values were calculated via likelihood ratio tests using the ‘afex’ package (Singmann et al., 2015).

**Figure 4:**
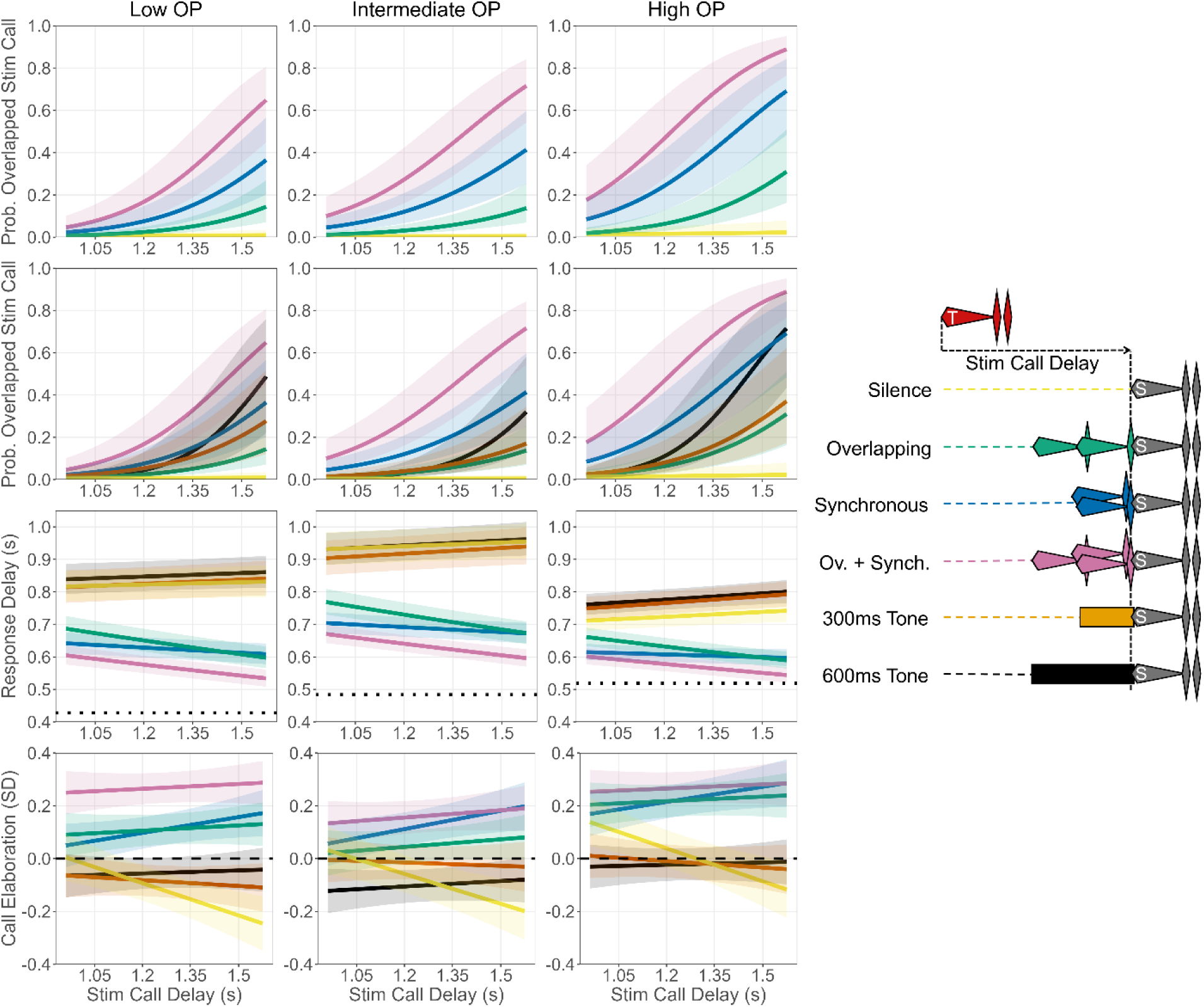
Visualization of the interaction between stimulus delay and acoustic motifs preceding stimulus calls in predicting various response call properties. Predictions and 95% confidence intervals; curve colors correspond to colors of motif depictions on the right. ***Top row***: Predicted probabilities that response calls overlapped stimulus calls (*Overlap_GLMM*), shown only for silence and conspecific call motifs, to reduce clutter. ***Second row***: Predicted probabilities that response calls overlapped stimulus calls (*Overlap_GLMM*), shown for all motifs. ***Third row***: Predicted response delays for response calls that did not overlap stimulus calls (*Response_Delay_LMM).* Dotted lines depict stimulus call durations. ***Fourth row***: Predicted call elaboration scores (standardized within males) relative to males’ mean elaboration levels (*Within_Male_Call_Elaboration_LMM*); dashed line at 0 indicates within-male mean elaboration score.

**Table 2:** Predictions and 95% CIs for control covariates from our models, across a representative range of predictor values. Predictions are for responses to stimulus calls preceded by the overlapping+synchronous motif at a stimulus delay of 1.5s (i.e., highly arousing conditions), and predictions are averaged across all stimulus call IDs. Grayed cells highlight statistically significant relationships; see Appendix for p-values.

| Predictor Variable: | Model Name: |  |  |  |
| --- | --- | --- | --- | --- |
|  | *Overlap_GLMM* | *Response_Delay_LMM* | *Within_Male_Call_Elaboration_LMM* | *Between_Male_Call_Elaboration_LMM* |
|  | Response Variable: |  |  |  |
|  | Overlap Probability | Response Delay (s) | Response Call Elaboration Within-Males (SD) | Response Call Elaboration Between-Males (score) |
| Body weight | **1.3g:** 0.33 [0.15, 0.59] | **1.3g:** 0.58 [0.54, 0.61] | - | **1.3g:** 2.4 [2.05, 2.82] |
|  | **1.7g:** 0.7 [0.53, 0.83] | **1.7g:** 0.56 [0.54, 0.59] | - | **1.7g:** 2.65 [2.41, 2.92] |
|  | **2.1g:** 0.92 [0.79, 0.97] | **2.1g:** 0.55 [0.51, 0.59] | - | **2.1g:** 2.93 [2.47, 3.47] |
| Water temperature | **26.5C:** 0.39 [0.12, 0.76] | **26.5C:** 0.59 [0.54, 0.64] | - | **26.5C:** 2.53 [1.98, 3.22] |
|  | **27.5C:** 0.66 [0.48, 0.81] | **27.5C:** 0.57 [0.54, 0.59] | - | **27.5C:** 2.63 [2.39, 2.9] |
|  | **28.5C:** 0.86 [0.6, 0.96] | **28.5C:** 0.55 [0.5, 0.59] | - | **28.5C:** 2.74 [2.22, 3.37] |
| Response call onset time | **5 minutes:** 0.75 [0.59, 0.86] | **5 minutes:** 0.57 [0.54, 0.49] | **5 minutes:** 0.16 [0.07, 0.25] | **5 minutes:** 2.52 [2.28, 2.8] |
|  | **17.5 minutes:** 0.68 [0.5, 0.82] | **17.5 minutes:** 0.57 [0.54, 0.49] | **17.5 minutes:** 0.25 [0.2, 0.31] | **17.5 minutes:** 2.65 [2.41, 2.91] |
|  | **30 minutes:** 0.61 [0.41, 0.78] | **30 minutes:** 0.56 [0.54, 0.49] | **30 minutes:** 0.35 [0.26, 0.45] | **30 minutes:** 2.78 [2.5, 3.09] |
| Trigger call elaboration | **-1SD:** 0.65 [0.46, 0.8] | **-1SD:** 0.57 [0.55, 0.6] | **-1SD:** -0.34 [-0.4, -0.28] | - |
|  | **mean:** 0.69 [0.51, 0.82] | **mean:** 0.57 [0.54, 0.59] | **mean:** 0.29 [0.23, 0.35] | - |
|  | **+1SD:** 0.72 [0.55, 0.85] | **+1SD:** 0.56 [0.54, 0.58] | **+1SD:** 0.92 [0.86, 0.98] | - |

#### Overlap_GLMM

Overlap probability was influenced by significant interactions between preceding motif and stimulus delay (p < 0.001), and preceding motif and stimulus call ID (p < 0.001), though the interaction between stimulus delay and stimulus call ID was non-significant (p = 0.13). All stimulus call IDs were essentially never overlapped at any stimulus delay when preceded by silence (predicted overlap probabilities [95% CI] when preceded by silence at 1.5s stimulus delay: low OP, 0.01 [0.0, 0.03]; intermediate OP, 0.01 [0.0, 0.02]; high OP, 0.02 [0.01, 0.07]). However, overlap probabilities increased when stimulus calls were preceded by our acoustic motifs, with the overlapping+synchronous motif inducing the highest probability of overlap (Figure 4). Additionally, overlap probabilities increased, and differences among motifs widened, as stimulus calls were presented at later stimulus delays. In contexts promoting overlap, the same hierarchy of stimulus call overlap probabilities observed in a live chorusing context was reproduced (predicted overlap probabilities [95% CI] when stimulus calls preceded by overlapping+synchronous motif at 1.5s stimulus delay: low OP, 0.54 [0.35, 0.72]; intermediate OP, 0.63 [0.46, 0.78]; high OP, 0.84 [0.68, 0.92]). 1000Hz tone motifs induced similar overlap probabilities to certain motifs composed of conspecific calls, suggesting generic inhibition by acoustic stimulation is an important driver of overlapping responses. Higher body weight and higher trigger call elaboration scores significantly increased overlap probabilities, whereas males became significantly less likely to overlap stimulus calls as trials wore on (Table 2). Though our fixed effects explained a good proportion of variation (marginal R^2^ = 0.3), most was explained by our random effects (conditional R^2^ = 0.78). The random intercept showed the highest variance (8.89), suggesting males differ widely in their baseline propensities to overlap rivals’ calls.

Durations of our different stimulus calls correlated positively with their probabilities of being overlapped (*Overlap_GLMM*). This gave us pause, as our working theory was that overlap is driven primarily by frequency and amplitude patterns throughout calls (Larter & Ryan, 2024b, 2024c), rather than generic features such as duration. However, a detailed look at overlap patterns reveals that the importance of duration arises indirectly due to increased call durations stemming primarily from additional chuck notes. Onsets of overlapping response calls occurred at several peaks during stimulus calls; most often during the latter portions of whines in accordance with our theory, but also tightly clustered during secondary chuck notes of multi-chuck stimulus calls (see intermediate OP and high OP in Figure 5). This primarily occurred when multi-chuck stimulus calls were preceded by motifs composed of conspecific calls which induced the shortest response delays (Figure 4, discussed below), rather than motifs composed of tones which induced long response delays. This suggests that this latter peak is generated by short-latency calls being triggered by the offsets of initial chucks. Thus, initial chucks may commonly trigger calls, but it is only when coupled with stimulation patterns that sufficiently shorten response delays that this results in this form of overlap.

**Figure 5.**
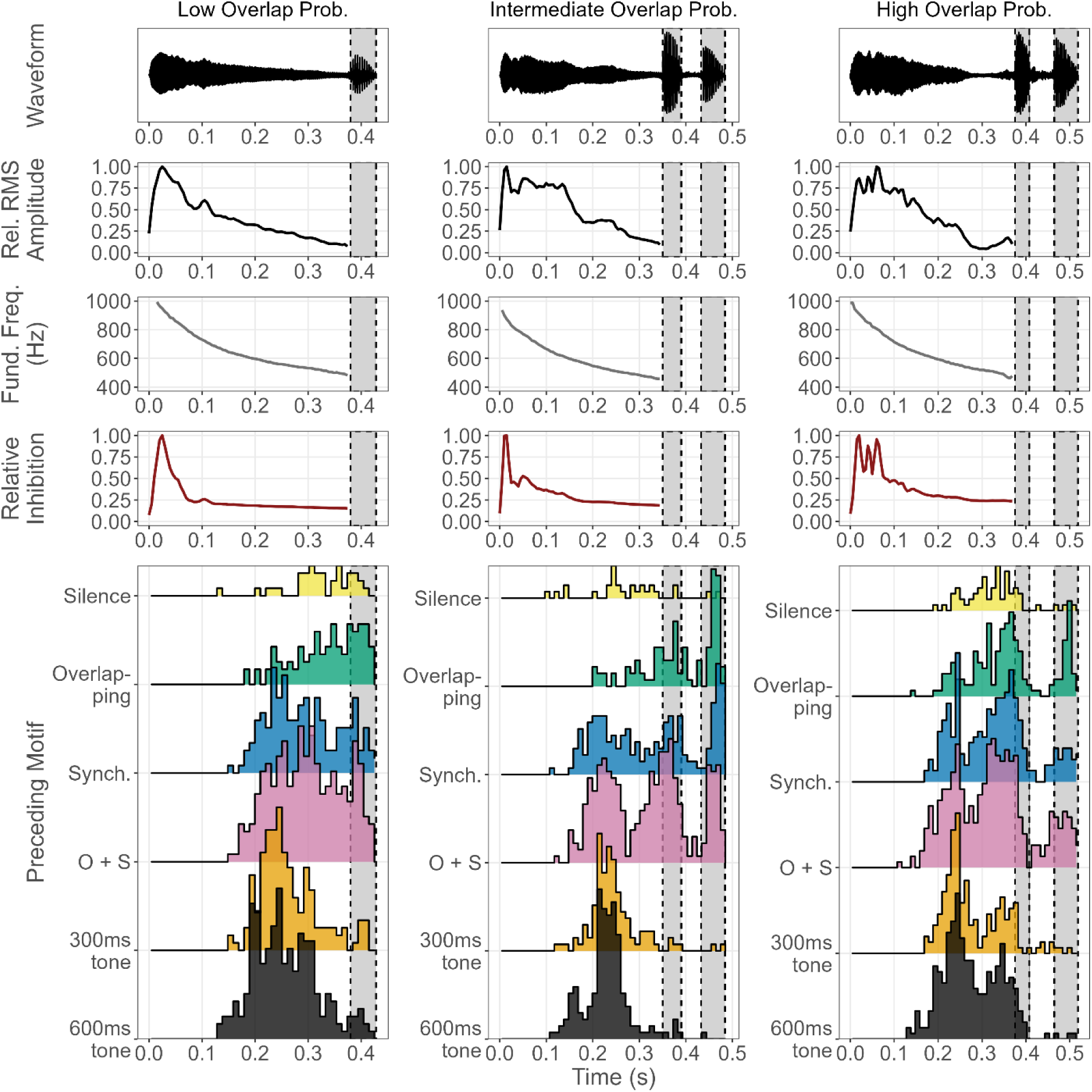
***Top***: Waveforms, relative RMS amplitude trajectory, fundamental frequency trajectory, and relative inhibition trajectories by stimulus call ID. Relative inhibition trajectories were obtained by projecting simultaneous amplitude and fundamental frequency trajectories into a male sensory system model generated in a calling context (Larter & Ryan, 2024b; illustrated in Figure 2 of Larter and Ryan, 2025; see SI7 for details). ***Bottom***: Ridgeline histograms (’ggridges’ R package: Wilke, 2018) showing when onsets of overlapping response calls occurred throughout stimulus calls when preceded by different motifs (10ms bins). Many calls occurred during the latter parts of whines as has been discussed previously (Larter & Ryan, 2024a, 2024c), but large, compact, peaks during secondary chucks are evident for intermediate and high OP calls; these calls were presumably triggered by offsets of initial chucks. Multiple peaks are also evident in the latter parts of whines of intermediate and high OP calls, and seem to arise due to inter-male variation in the timing of overlapping calls (see SI8 for visualizations and discussion). See SI9 for similar histograms for all responses (overlapping and non-overlapping).

To disentangle this form of overlap from the stereotyped overlap arising when species-typical whine amplitude and frequency trajectories trigger response calls (Larter & Ryan, 2024b), post-hoc we ran an identical model to *Overlap_GLMM*. Except, here, we only considered response calls to be overlapping stimulus calls if their onsets occurred prior to the ends of the initial chucks of intermediate OP and high OP calls (***Restricted_Overlap_GLMM***). Results differed somewhat from *Overlap_GLMM*, with intermediate OP having a slightly lower overlap probability than low OP (though these were essentially equivalent), but with high OP still having the highest overlap probability overall (predicted overlap probabilities [95% CI] when stimulus calls preceded by overlapping+synchronous motif at 1.5s stimulus delay: low OP, 0.53 [0.33, 0.72]; intermediate OP, 0.49 [0.31, 0.67]; high OP, 0.7 [0.49, 0.85]). See SI6 for visualizations.

#### Response_Delay_LMM

For responses that did not overlap stimulus calls, response delays were influenced by significant interactions between preceding motif and stimulus delay (p < 0.001), and preceding motif and stimulus call ID (p < 0.001), though the interaction between stimulus delay and stimulus call ID was non-significant (p = 0.18). The effects of motifs separated into two clear clusters (Figure 4); stimulus calls preceded by silence and 1000Hz tone motifs induced similarly long response delays, which increased slightly as stimulus calls were encountered at greater stimulus delays. Conversely, motifs composed of conspecific calls induced shorter response delays that tended to decrease somewhat as stimulus calls were encountered at greater stimulus delays, and response delay distributions (see SI8) exhibited pronounced peaks ∼60ms after the offsets of stimulus calls (near lower reaction-time limits for this species: Larter & Ryan, 2024b). Stimulus call ID influenced average response delays in ways not in accordance with the hierarchy of stimulus call durations (low OP < Intermediate OP < high OP) (predicted response delays (s) [95% CI] averaged across all preceding motifs at 1.5s stimulus delay: low OP, 0.7 [0.67, 0.74]; intermediate OP, 0.79 [0.75, 0.83]; high OP, 0.67 [0.65, 0.7]), suggesting the stimulus calls themselves aroused males to different degrees. Our control variables had only negligible and/or non-significant effects (Table 2). Here, most variation was explained by our fixed effects (marginal R^2^ = 0.39, conditional R^2^ = 0.73).

#### Within_Male_Call_Elaboration_LMM

Response call elaboration, standardized to reflect within-male changes relative to typical call elaboration levels, was influenced by significant interactions between preceding motif and stimulus delay (p < 0.001), and preceding motif and stimulus call ID (p = 0.004), though the interaction between stimulus delay and stimulus call ID was non-significant (p = 0.4). Here again, the same two clusters are evident as for *Response_Delay_LMM* (Figure 4). When males produced trigger calls at their average level of elaboration, motifs composed of silence or 1000Hz tones tended to promote maintenance of, or a slight decrease of, relative call elaboration levels in response calls. For stimulus calls preceded by silence, we also see a pronounced temporal effect, with longer stimulus delays inducing greater reductions in response call elaboration. Conversely, motifs composed of conspecific calls tended to increase response call elaboration relative to trigger calls, and this effect increased somewhat at greater stimulus delays. Males also tended to increase call elaboration throughout trials, and greater trigger call elaboration scores predicted greater response call elaboration (Table 2), demonstrating the inertia over time in call elaboration levels seen previously (Bernal et al., 2009). In fact, response call elaboration score was almost entirely predicted by the elaboration score of the trigger call immediately preceding it; marginal R^2^ for the full model was 0.47 (conditional R^2^ = 0.52), and removing trigger call elaboration score as a predictor reduced the marginal R^2^ to 0.04, whereas removing preceding motif reduced it to 0.45.

#### Between_Male_Call_Elaboration_LMM

In agreement with previous work (Bernal et al., 2007), absolute response call elaboration levels were not significantly influenced by male body weight or water temperature, though they were positively associated (Table 2). In fact, almost all variation in this model was explained by the random effects (marginal R^2^ = 0.06; conditional R^2^ = 0.76). Thus, though males differ consistently in call elaboration patterns, potential constraints remain a mystery (Larter & Ryan, 2024a). See SI10 for call elaboration score distributions.

## Discussion

Many taxa form collective behavioral aggregations, and optimal interaction patterns often differ across social contexts. Sensory scenes arising in dense social environments are complex, selecting for mechanisms allowing individuals to distill informative cues from these scenes to guide appropriate response adjustments. Here, we used automated playback to investigate how conspecific stimulation patterns typical of different chorusing environments interacted with endogenous responsiveness rhythms to guide the socially-mediated flexibility in interaction patterns observed in live túngara frog choruses. Our results revealed that this flexibility is driven on a call-by-call basis by a multifaceted cue to social context; the character of short-term conspecific stimulation patterns experienced prior to each call, and how the timing of this stimulation interacts with cyclical responsiveness changes unfolding throughout call cycles. Overall, our results demonstrate that seemingly inactive periods within behavioral rhythms provide valuable repeated assessment windows for fine-tuning upcoming responses, and that responsiveness cycles serve as important temporal filters for mapping conspecific stimulation patterns to behavioral outputs.

### Temporal Responses to Isolated Conspecific Calls

Phase-response curves (PRCs) have been characterized extensively in chorusing insects (Greenfield & Roizen, 1993; Sismondo, 1990; Walker, 1969). These can be complex and discontinuous functions, with males of some species exhibiting varied temporal responses to rivals’ calls encountered at different points throughout their call cycles. Fewer frog PRCs are available, but those available show simpler continuous functions driven by consistent short-latency responses to rival’s calls encountered at all delays beyond the refractory period (Lemon & Struger, 1980; Loftus-Hills, 1974; Larter and Ryan 2025). Túngara frogs exhibit this same pattern, with our response delay curve remaining flat for all stimulus delays beyond the initial refractory period, and with the response period curve showing a corresponding 1:1 linear increase (Figure 3).

Though we did not systematically sample stimulus delays that overlapped male trigger calls (< 0.45s), opportunistic data from a subset of males suggest that stimulus calls at these delays cannot elicit the short-latency responses seen at later delays (illustrating the refractory period: Narins, 1982). However, the decreasing left arm of our curves suggests that stimulus calls encountered during the refractory period still shorten the current call period somewhat, and do so to greater degrees at later stimulus delays throughout the refractory period (also seen in Pseudacris crucifer: Lemon & Struger, 1980). This suggests that these calls are perceived to some degree and arouse males such that the next endogenously triggered call occurs sooner than it otherwise would have. The increasing effect at later delays likely occurs because larger portions of later overlapping stimulus calls are perceivable protruding beyond the ends of males’ own trigger calls.

### Temporal Responses in the Context of Conspecific Stimulation Patterns

Dyadic interactions represent but one of countless social contexts in which call-timing mechanisms must function, yet this is typically the sole context in which they are investigated. However, the simple and consistent response patterns exhibited by túngara frogs in a dyadic context (Figure 3) belie the true versatility of their call-timing mechanism. This versatility was revealed only when we presented stimulus calls to males preceded by acoustic motifs that mimicked conspecific stimulation patterns typically encountered in different social contexts (we discuss the effects of tonal motifs in a later section). Isolated stimulus calls were essentially never overlapped at any stimulus delay. However, when stimulus calls were preceded by motifs consisting of combinations of overlapping and synchronous conspecific calls (interaction types common in larger choruses: Larter & Ryan, 2024a), males altered their call-timing responses; by increasing their probabilities of overlapping stimulus calls in the stereotyped fashion seen in this species, and decreasing the latencies of their response calls relative to call-triggers (Figure 4; SI8). This was most pronounced for the motif featuring both overlapping and synchronous conspecific calls (overlapping+synchronous), which exhibited both the greatest duration and highest peak and average amplitude of the conspecific motifs used here.

Claiming that stereotyped overlapping responses can only be elicited from males when their rivals are already engaging in them would be circular reasoning. Thus, it is almost certainly not that the specific conspecific interaction motifs used here are crucial for generating the observed correlation between overlap prevalence and chorus size. Rather, it is likely that experiencing intense conspecific stimulation more generally during call cycles arouses males in larger choruses and increases their probability of overlapping. In larger choruses, chorus duty cycles increase (Larter & Ryan, 2024a), and males increase call amplitude (Larter et al., 2022) and call elaboration (Bernal et al., 2007; Larter & Ryan, 2024a). Consequently, conspecific stimulation becomes more intense and more constant as choruses grow, with overlapping and synchronous calls further exaggerating these effects.

Furthermore, we observed a salient interaction between motifs and males’ endogenous responsiveness rhythms in influencing overlap probabilities and response latencies, with the escalatory effects of conspecific motifs becoming exaggerated at later stimulus delays (Figure 4). We played all stimulus call/stimulus delay/motif combinations to males in a random order within the same playback session, interspersed with standardized alternation interactions to reduce potential carry-over effects. The varied effects of different stimulus combinations therefore indicate that responses to rivals’ calls arise as the parameters of male call-timing mechanisms are shaped anew each call cycle, by a complex interaction between their cyclical endogenous responsiveness rhythms and the conspecific stimulation patterns they sample prior to each impending call.

Regarding the stereotyped overlapping responses observed in this species (Larter & Ryan, 2024a, 2024c), the crucial parameter appears to be how permissive male gap-detection mechanisms are regarding the magnitude of a release from inhibition that is sufficient to trigger a call. The moderate release from inhibition shortly following rivals’ whine onsets becomes an increasingly salient call-trigger in larger choruses due to male gap-detection mechanisms becoming increasingly permissive within these social contexts, leading to the observed correlation between overlap prevalence and chorus density (Larter & Ryan, 2024b). Our results here suggest that this correlation arises as male call-timing mechanisms tailor response patterns to social contexts on a call-by-call basis, by altering this crucial permissiveness parameter each call cycle in response to an emergent, multifaceted, cue generated by rivals in different social environments; that, as choruses grow larger, males experience increasingly intense conspecific stimulation patterns between calls, and increasingly experience them throughout the latter reaches of their call cycles. The responses that ultimately terminate each call cycle then arise as this freshly altered call-timing mechanism interacts with the acoustic properties of rivals’ calls (and emergent features of acoustic scenes: Larter & Ryan, 2024c) encountered subsequent to these alterations. Hence the varied responses to different stimulus call IDs when preceded by the same motifs at the same delays. That, when using the same overlap metric (*Overlap_GLMM*), we were able to experimentally recover the same hierarchy of overlap probabilities for our stimulus calls as was observed in a live 6-male chorus (Larter & Ryan, 2024c) provides strong support for this proposed mechanism.

### Túngara Frog Call-Timing Mechanisms

Our emerging picture of the functioning of túngara frog call-timing mechanisms reveals properties that are likely advantageous across the varied range of social environments in which males can find themselves. The switch from alternation in smaller choruses (≤ 3 males) to increasingly prevalent stereotyped call overlap in larger choruses is beneficial, as overlapping rivals’ calls in this stereotyped way imposes the lowest attractiveness costs relative to all other potential forms of overlap (Larter & Ryan, 2024a). Thus, becoming more generally permissive of overlap when it becomes unavoidable in larger choruses allows males to maintain high call rates, while the specific sensory tuning of gap-detection mechanisms strongly prioritizes overlap of this least costly form (Larter & Ryan, 2024b). Similarly, reduced response delays in crowded choruses allow males to most effectively insert calls into short-lived lulls in background chorus noise (Larter & Ryan, 2024a).

Furthermore, chorus noise is temporally-heterogenous, and that response patterns are updated on a call-by-call basis allows the tradeoff between calling at high rates and calling at times of reduced interference to be continuously calibrated to short-term chorus dynamics (Larter and Ryan, 2025). Males can immediately become permissive of overlapping rivals’ calls in this least costly way when intense, continuous, inter-call stimulation patterns indicate that encountering silent gaps within the time horizon of the current call period is unlikely. Yet, they can immediately switch back to prioritizing calling within silent gaps again when sparser inter-call stimulation patterns suggest lengthy silences may be forthcoming (non-overlapping calls are most preferred by females overall: Legett et al., 2019). Indeed, overlap and alternation both occur in larger choruses (Larter & Ryan, 2024a). Thus, our results demonstrate that inactive periods during behavioral cycles afford valuable repeated opportunities to sample dynamic social scenes, such that each impending response can be fine-tuned in light of current collective behavior patterns (Ariel et al., 2014). The revealed malleability of túngara frog call-timing mechanisms in response to short-term stimulation patterns contrasts with traditional models that depict call-timing mechanisms as essentially rigid across social/acoustic contexts (Larter & Ryan, 2025).

### Call Elaboration in the Context of Conspecific Stimulation Patterns

In agreement with the positive correlation between chorus size and call elaboration observed previously in live túngara frog choruses (Bernal et al., 2007; Larter & Ryan, 2024a), call elaboration was also increased somewhat by conspecific motifs typical of larger choruses. We observed the same general motif arousal hierarchy as was seen in temporal contexts (overlapping+synchronous motifs being the most arousing), and similar (though less pronounced) increases in this effect as stimulus calls were encountered at later delays. Conversely, when motifs were preceded by silence, we saw a tendency to reduce call elaboration, with this reduction becoming starker as stimulus calls were encountered at later delays (yellow lines in lower plot of Figure 4). Túngara frog calling strategies have been strongly influenced by the behavior of predatory bats and parasitic flies that eavesdrop on male calls to locate them as prey, and show similar preferences for complex calls as do females (Bernal et al., 2006; Tuttle & Ryan, 1981). Consequently, males rapidly cease calling when their nearby rivals cease, as this suggests these rivals may have detected an incoming eavesdropper, and because males still calling when others go silent will face higher, less diluted, risks (Dapper et al., 2011). The apparent rapid decay of arousal over the course of call cycles exhibiting extended silences is likely part of this defensive response.

Overall though, response call elaboration was primarily driven by the elaboration of the trigger call preceding it. Thus, in contrast to overlap probability and response latencies which are primarily responsive to short-term inter-call stimulation patterns, call elaboration is subject to a high degree of inertia over longer stretches of calling (as shown previously: Bernal et al., 2009), and is only influenced slightly by shorter-term stimulation patterns. This suggests that call elaboration and temporal aspects of responses are controlled by different neural circuits that integrate sensory data over different time horizons. However, that motifs did influence response call elaboration to some degree, and that males calling more elaborately exhibited slightly higher overlap probabilities (Table 2), suggests these circuits partially intersect and interact.

### Similar and Divergent Influences of Conspecific and Tonal Motifs

Though conspecific motifs had broadly analogous escalatory effects on all aspects of call-timing responses we investigated, effects of tonal motifs showed interesting differences across response properties. Tonal motifs induced longer response delays and slight reductions in call elaboration, thus having similar effects to silence for these call properties (Figure 4; SI8). However, tonal motifs had a similarly positive effect to certain conspecific motifs (e.g., overlapping call motifs, and synchronous call motifs) on overlap probabilities. This suggests that, though conspecific stimulation influences all of these response properties in similarly escalatory ways, it may be different aspects of this stimulation that are salient. For instance, stimulation by more specific conspecific patterns (e.g., species-typical amplitude and frequency trajectories: Ryan, 1983; Walkowiak, 1988) may be necessary to induce shorter response delays and increased call elaboration. Conversely, more generic effects of conspecific stimulation patterns, such as inhibition by high-amplitude acoustic stimulation generally, may be the salient feature driving overlap probabilities, hence our tonal motifs eliciting similar effects.

It may be that broader-tuning of the elements of call-timing mechanisms that calibrate gap-detection behavior to background interference levels is beneficial, as this would allow callers to be similarly responsive to patterns of background noise arising from other species in mixed-species choruses, and abiotic noise, which also interfere with call transmission (Vélez et al., 2013). Whereas more specificity in the tuning of elements dealing with, e.g. call elaboration, would ensure males only incur the risks/costs of calling elaborately when conspecific competition is high and elaboration will be beneficial, and eavesdropper risks are diluted by calling rivals (Bernal et al., 2006; Ryan et al., 1981; Tuttle & Ryan, 1981). That different features of conspecific stimulation patterns seem to drive different properties of responses emphasizes the complexity of the interactions between the neural and sensory processes governing response flexibility across several axes (Walkowiak, 1988), and the multi-dimensional, ever-changing, sensory scenes that guide responses (Hein, 2022; Williams et al., 2023).

## Conclusions

Our findings demonstrate that túngara frogs sample emergent patterns in conspecific scenes during the inactive periods preceding each call, and alter their interaction mechanisms to facilitate beneficial responses given this immediate context. Furthermore, salient features of conspecific scenes for guiding these adjustments are multifaceted, involving both the character of stimulation perceived, and when it is perceived relative to males’ endogenous responsiveness rhythms. Pieces of this picture are evident in other collective behavioral contexts, suggesting these results have broader relevance beyond chorusing. For instance, behavioral adjustments in response to broader emergent properties of conspecific sensory scenes facilitate collective hunting in spiders (Chiara et al., 2022) and collective locomotion in zebrafish (Harpaz et al., 2021). Additionally, alternating phases of sampling collective dynamics, and responding in light of these dynamics, underpin group cohesion during collective locomotion in locust and zebrafish groups (Ariel et al., 2014; Harpaz et al., 2017). Thus, cues arising as emergent patterns in conspecific sensory scenes are filtered through the temporal prisms of individual responsiveness rhythms likely have widespread importance for guiding appropriate interaction patterns within dynamic behavioral collectives.

## Acknowledgements

We thank the editors and reviewers, and members of the Ryan Lab, for helpful comments on the manuscript. We thank Gregg Cohen and the Smithsonian Tropical Research Institute for logistical support. We thank Matías Muñoz for providing Figure 1 code.

## Data Accessibility

Data and code are available at Dryad repository (Larter, Cushing, & Ryan, 2025).

## Appendix 1: GLMM Summaries

**Figure A1:**
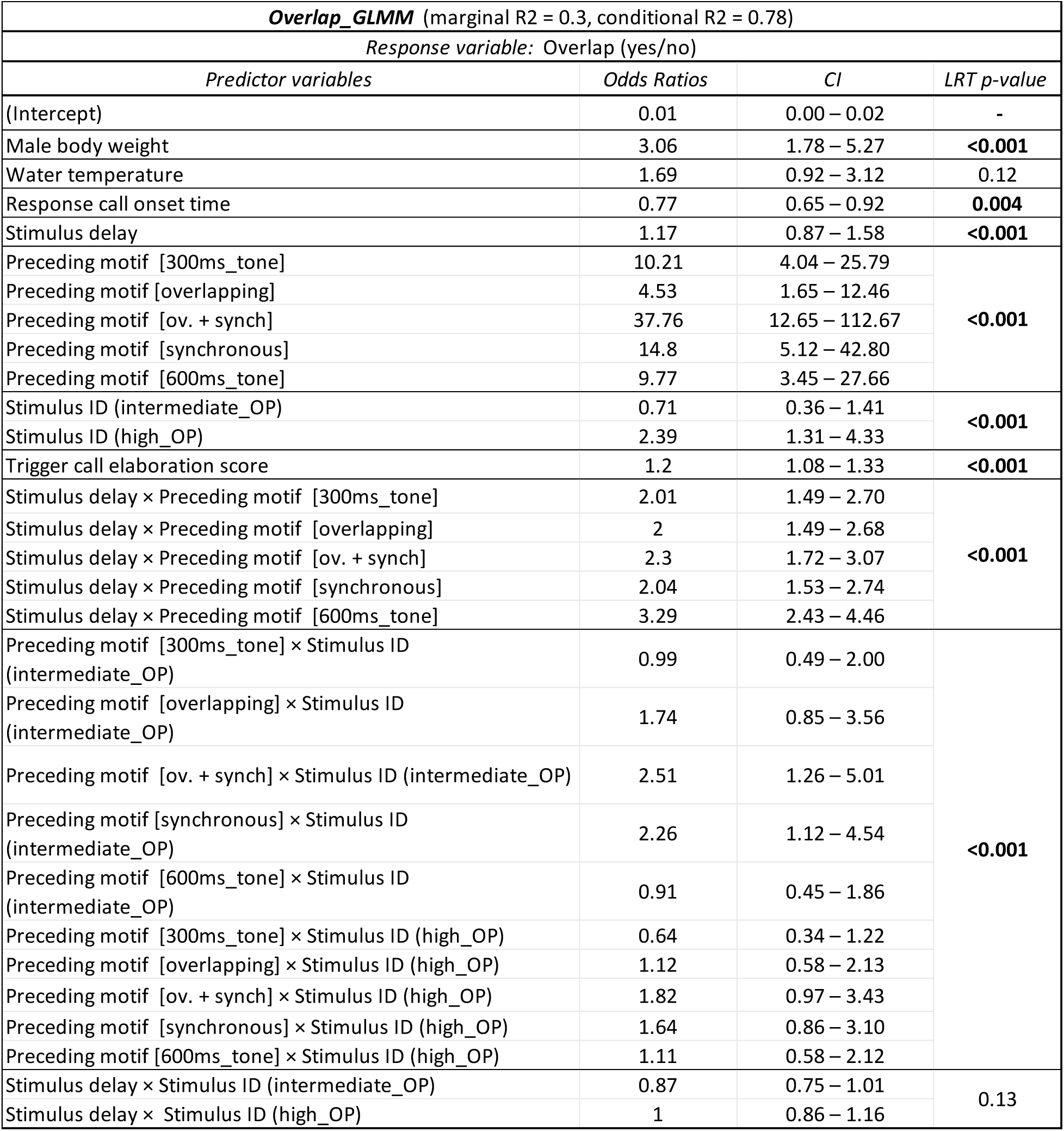
Overlap_GLMM Summary.

**Figure A2:**
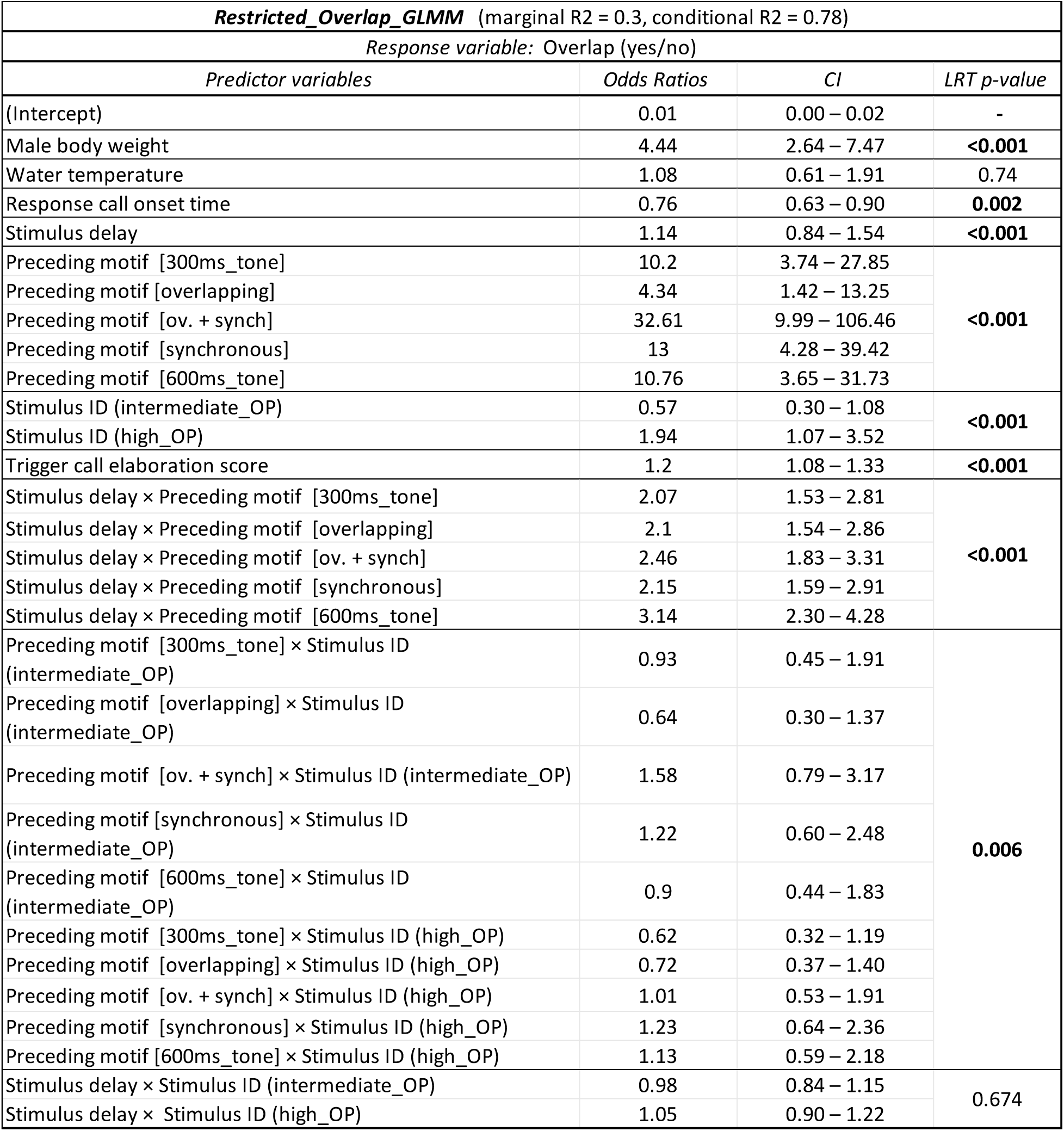
Restricted_Overlap_GLMM Summary.

**Figure A3:**
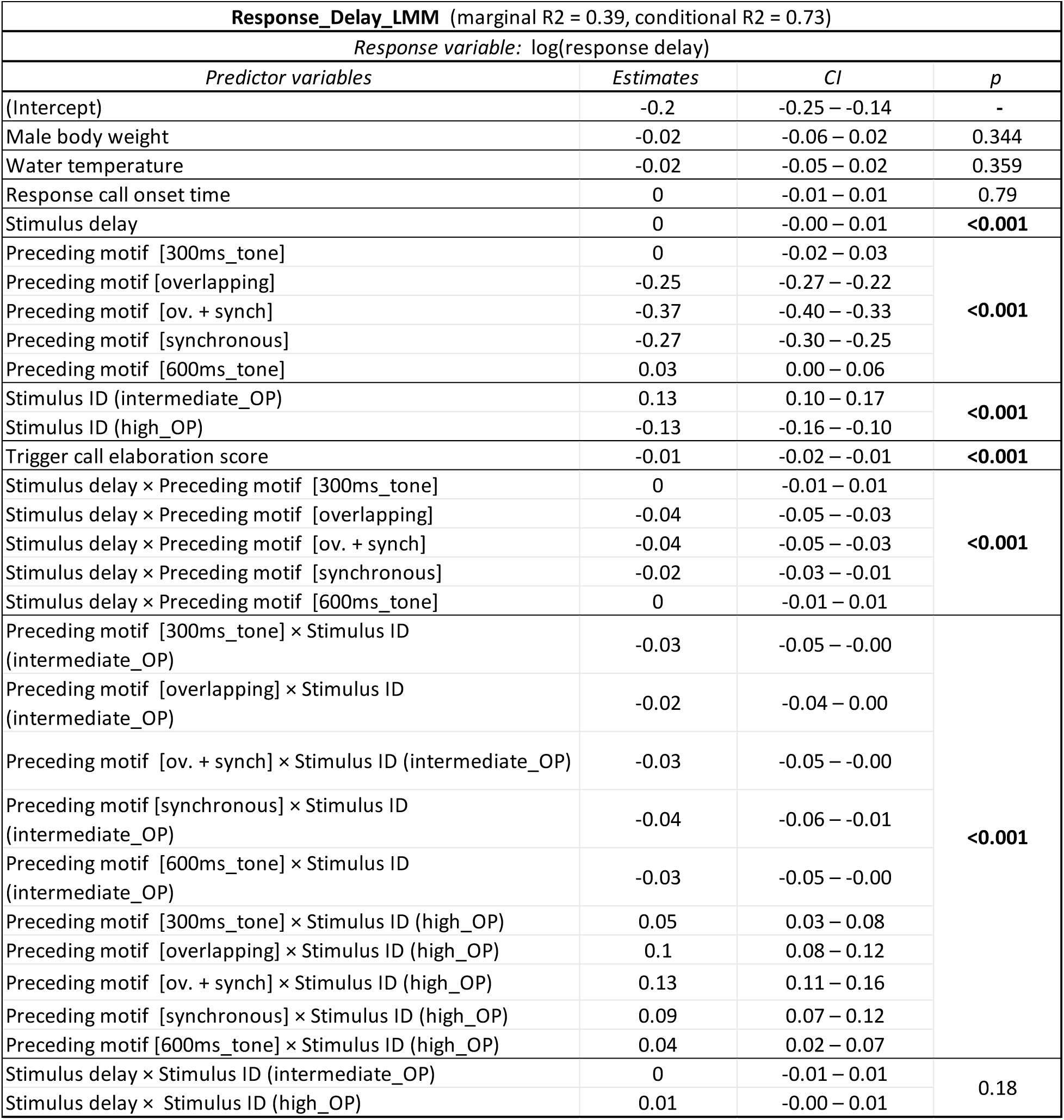
Response_Delay_LMM Summary.

**Figure A4:**
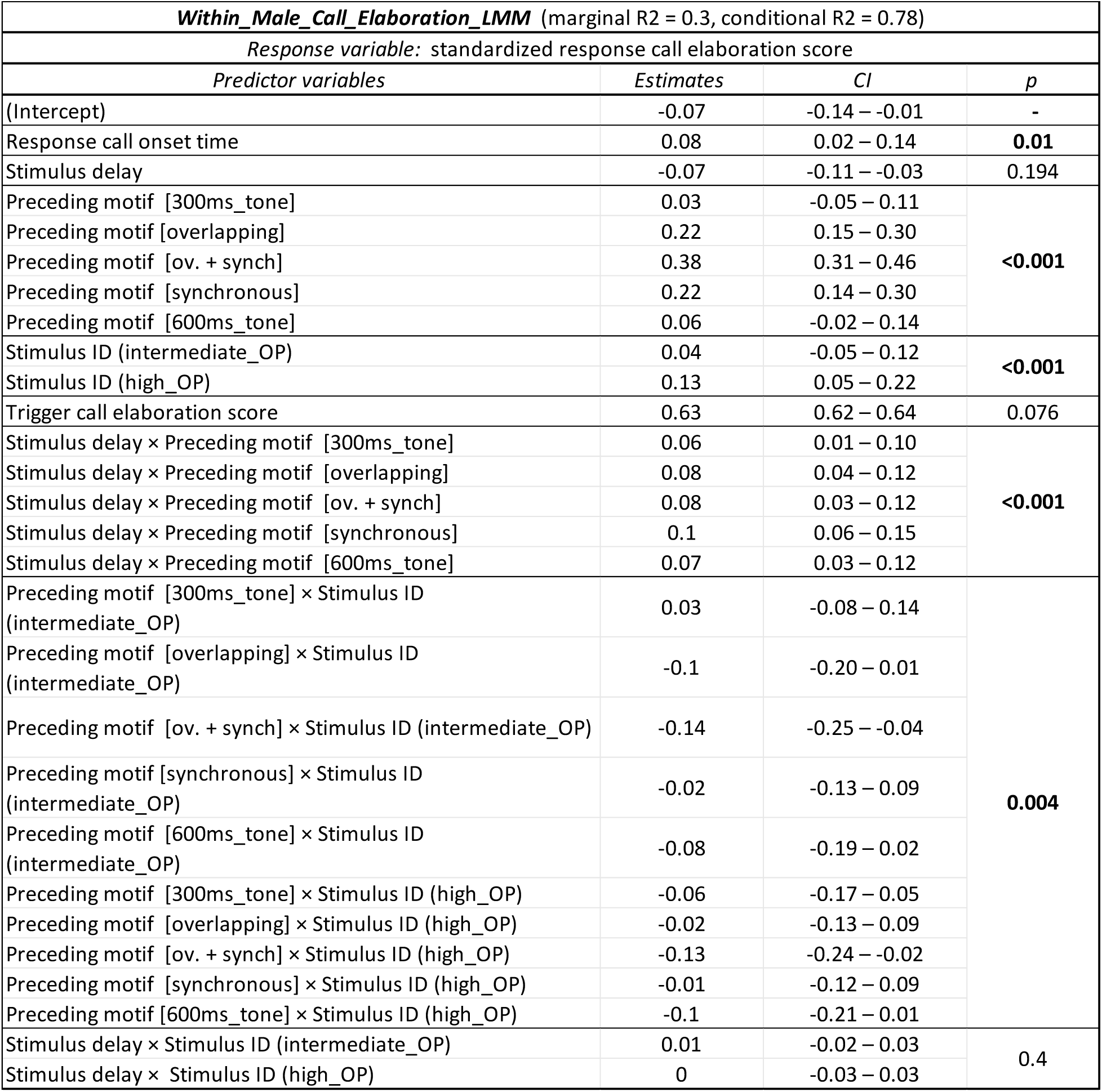
Within_Male_Call_Elaboration_GLMM Summary.

**Figure A5:**
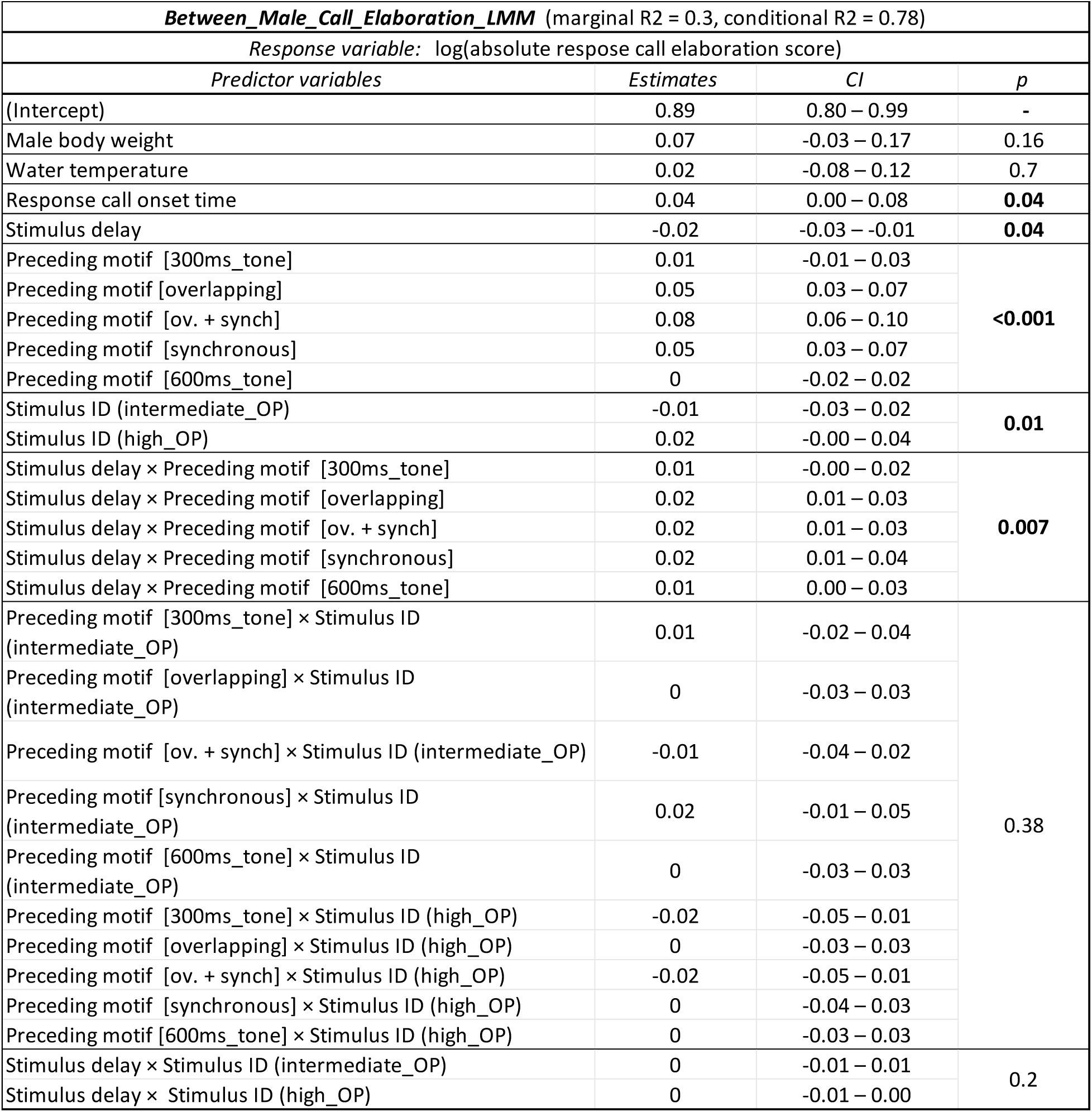
Between_Male_Call_Elaboration_GLMM Summary.

