## Supplementary material for "Cadences of the Collective: Conspecific Stimulation Patterns Interact with Endogenous Rhythms to Cue Socially Mediated Response Shifts": SI

#### SI1: Visualizations of Playback Motifs

Spectrograms (upper), waveform (lower), and power spectrum (right) of playback motifs described in-text preceding the intermediate OP stimulus call ('seewave' R package, window length 1024, overlap 90%). Brighter colors denote higher intensities.

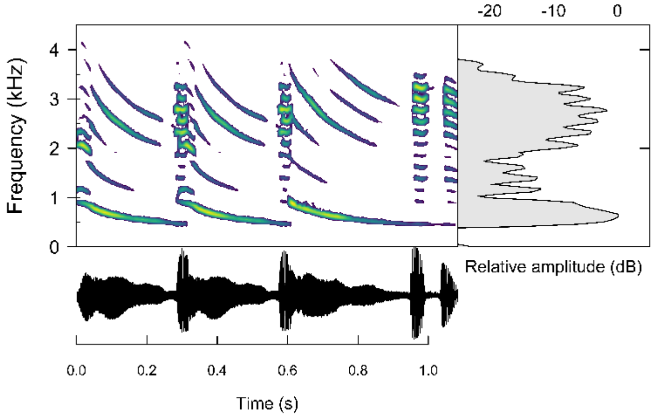


Figure SI1A: Overlapping motif

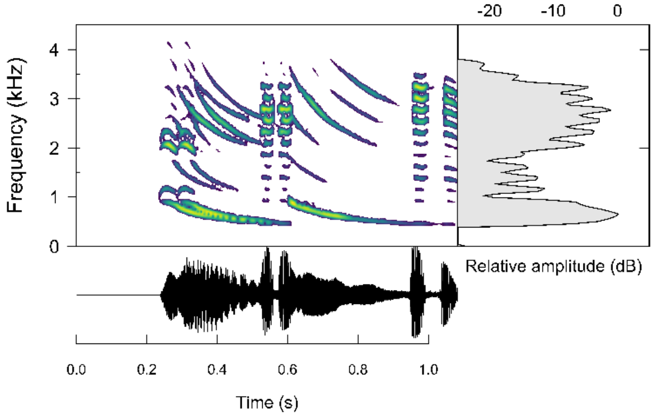


Figure SI1B: Synchronous motif

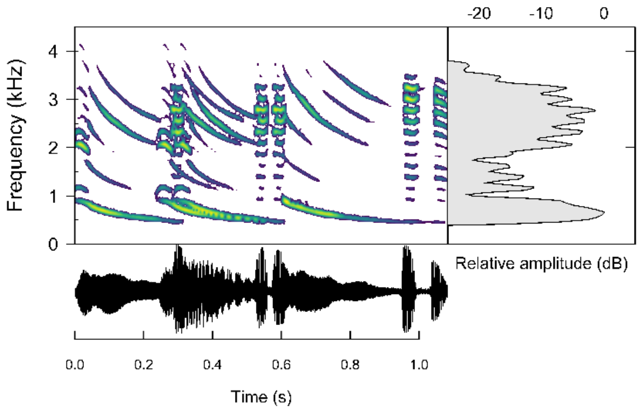


Figure SI1C: Synchronous+overlapping motif

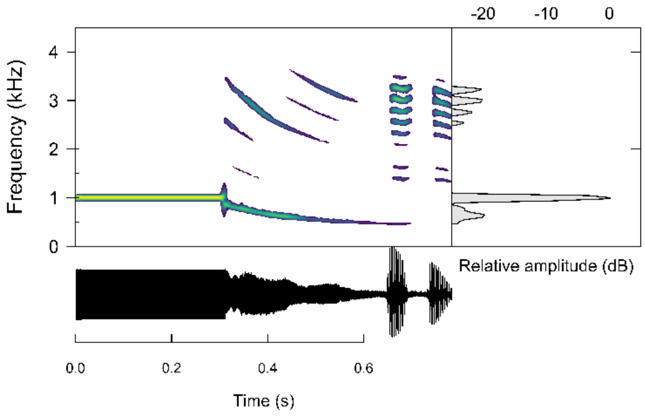


Figure SI1D: 300ms 1000Hz tone motif

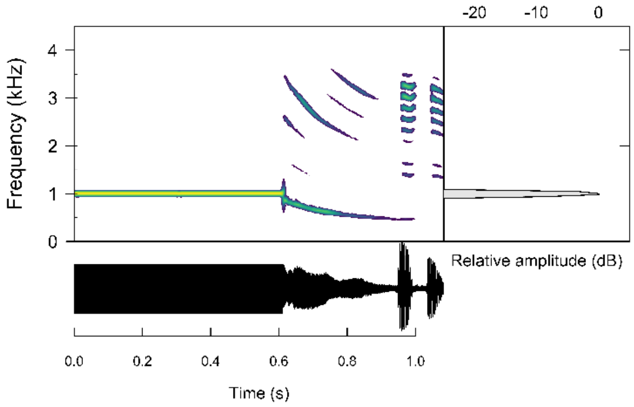


Figure SI1E: 600ms 1000Hz tone motif

SI2: Inter-male Variation in Response Delay Curves

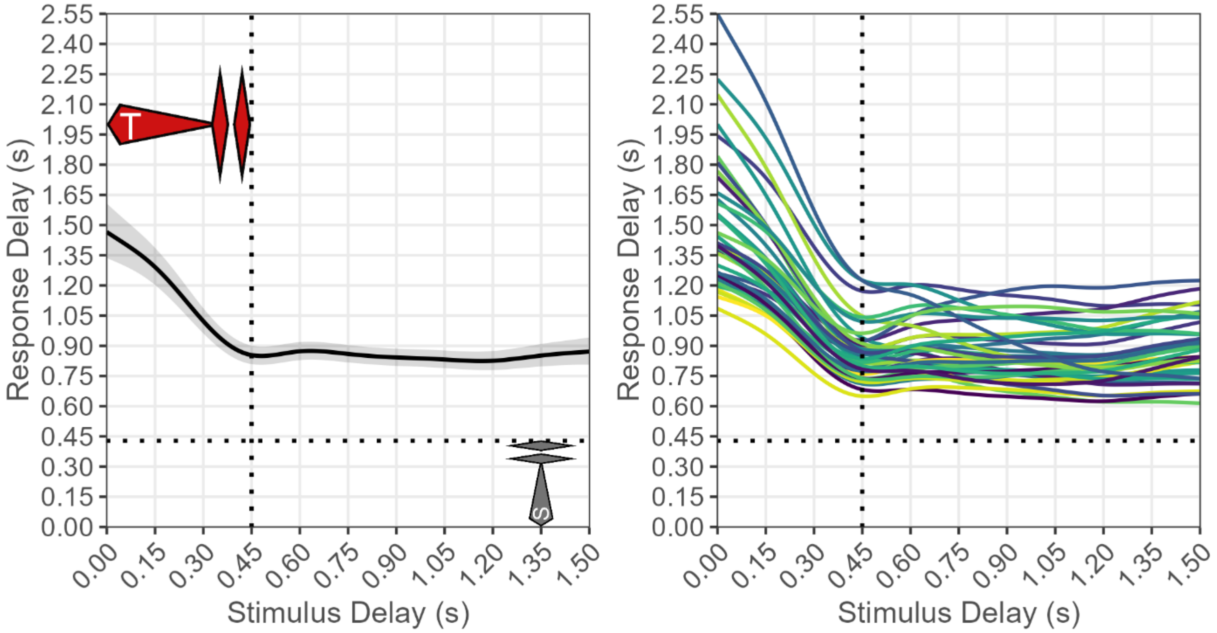


**Figure SI2**: **Left**) Response_Delay_GAMM results: Population average response delays to isolated low OP stimulus calls encountered at various stimulus delays. Plotting responses to a single stimulus call ID allows us to visualize responses relative to a specific stimulus call duration; low OP was chosen arbitrarily. Red call approximates a typical trigger call and the gray call represents the stimulus call; thus the vertical dotted line illustrates the duration of typical calls (Ryan & Rand, 2003) with playback delays to the left of this line representing stimuli that likely overlapped trigger calls to various degrees. Horizontal dashed line illustrates the duration of the low OP stimulus call. **Right**) Response delay curves for each individual male in our dataset. As can be seen, all males' curves follow a similar pattern, with most variation being in curve intercept.

SI3: Response Period Curve Model Results

| ***Response_Period_GAMM*** (R2 = 0.88) | | | | |
|---|---|---|---|---|
| *Linear Terms* | | | | |
| *Predictor variables* | *Estimates* | *CI* | | *p* |
| (Intercept) | 1.7 | 0.43 – 6.61 | | 0.45 |
| Response call onset time | 1 | 1.00 – 1.00 | | 0.76 |
| Trigger call elaboration score | 0.99 | 0.99 – 1 | | **0.002** |
| Stimulus call ID (intermediate OP) | 1.04 | 1.03 – 1.05 | | **<0.001** |
| Stimulus call ID (high OP) | 0.95 | 0.94 – 0.96 | | **<0.001** |
| Body weight | 0.91 | 0.84 – 0.99 | | **0.024** |
| Water temperature | 1 | 0.96 – 1.06 | | 0.85 |
| *Smooth Terms* | | | | |
| *Predictor variables* | *edf* | *Ref.df* | *F* | *p* |
| Smooth term (stimulus delay) | 8.65 | 8.94 | 128.74 | **<0.001** |
| Random intercept (male ID) | 32.33 | 36 | 142.29 | **<0.001** |
| Smooth term (stimulus delay, by male ID) | 110.5 | 386 | 19.43 | **<0.001** |
| Random Slope (response call onset time, by male ID) | 26.37 | 38 | 35.87 | **<0.001** |
| Random Slope (trigger call elaboration score, by male ID) | 21.88 | 38 | 2.11 | **<0.001** |
| Random slope (stimulus call ID, by male ID) | 58.66 | 112 | 2.25 | **<0.001** |

**Table SI3**: Results of *Response_Period_GAMM* (response period ~ stimulus delay). Model is described and results are visualized in-text. Predictor variables for GAMMs were not standardized prior to inclusion, with the exception of trigger call elaboration score which was standardized within males.

### SI4: Biased Sampling Among Males for Stimulus Delays that Overlapped Trigger Calls

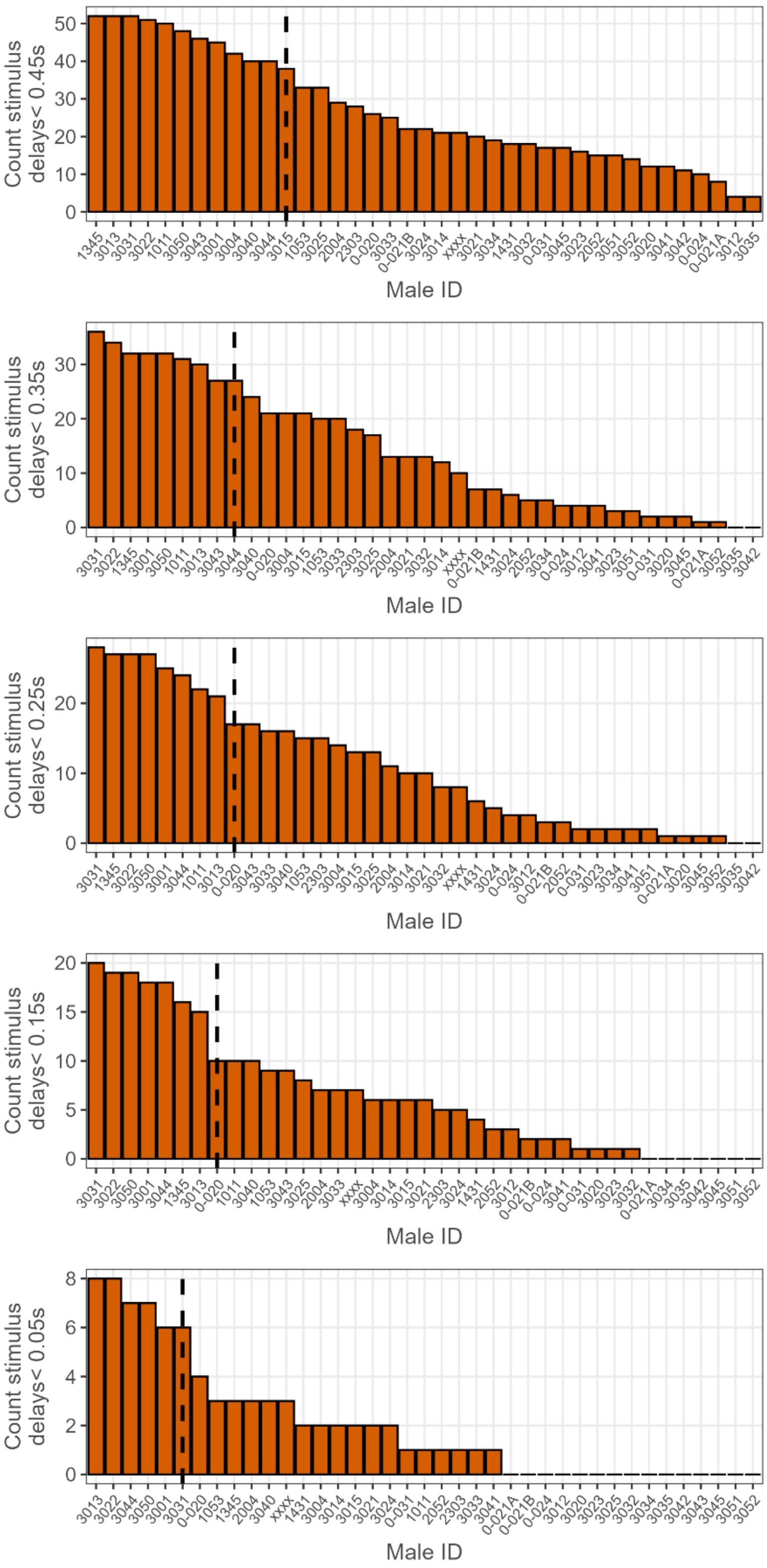


**Figure SI4**: Demonstration of the bias in our results for the shortest stimulus delays included in our response delay and response period curves. As we only obtained data from these shorter delays (<0.45s) opportunistically when males called spontaneously just prior to the onsets of stimulus calls at later delays (latest stimulus delay was 1.5s), responses at these delays are biased towards males who routinely exhibited call periods <1.5s, i.e. those with the shortest typical call periods. This bias is shown by these histograms, with the count of responses contributed by each male at stimulus delays <0.45s (or shorter) on the y axis, and males arranged on the x axis by counts of contributed responses at these delays. The vertical dotted line represents 50% of responses; i.e., the male bisected by the line and those with higher counts comprise 50% of the responses at these short delays. This demonstrates the bias towards a subset of males, and that this bias gets more extreme as we look at shorter and shorter stimulus delays. This bias explains the lower-than-expected intercept of these curves (lower than the expected call period), discussed in figure 3 caption in-text. Thus, if we had sampled responses to these shorter stimulus delays more systematically, we would likely see an initial arm of these curves more similar to the inferred red dotted line in figure 3.

### SI5: Random Effects Summaries for GLMMs and LMMs

Overlap_GLMM

| Group | RE Type | Associated Fixed Effect | # Levels | Variance | Std. Dev. |
|---|---|---|---|---|---|
| male_ID | Intercept | - | 39 | 8.89 | 2.98 |
| male_ID | Correlated random slope | Response_call_onset_time | 39 | 0.21 | 0.46 |
| male_ID | Correlated random slope | stimulus_call_OP_ordinal_facIntermediateOP | 39 | 0.63 | 0.8 |
| male_ID | Correlated random slope | stimulus_call_OP_ordinal_facHighOP | 39 | 0.39 | 0.63 |
| male_ID | Correlated random slope | Motif_preceding_stim_call300ms_tone | 39 | 2.42 | 1.55 |
| male_ID | Correlated random slope | Motif_preceding_stim_call600ms_ol_only | 39 | 4 | 2 |
| male_ID | Correlated random slope | Motif_preceding_stim_call600ms_ol_synch | 39 | 5.97 | 2.44 |
| male_ID | Correlated random slope | Motif_preceding_stim_call600ms_synch_only | 39 | 5.13 | 2.27 |
| male_ID | Correlated random slope | Motif_preceding_stim_call600ms_tone | 39 | 4.37 | 2.1 |
| male_ID | Uncorrelated random slope | Trigger_call_elaboration_score | 39 | 0.05 | 0.23 |
| male_ID | Uncorrelated random slope | Stimulus_delay | 39 | 0.14 | 0.37 |

Restricted_Overlap_GLMM

| Group | RE Type | Associated Fixed Effect | # Levels | Variance | Std. Dev. |
|---|---|---|---|---|---|
| male_ID | Intercept | - | 39 | 7.83 | 2.8 |
| male_ID | Correlated random slope | Response_call_onset_time | 39 | 0.2 | 0.45 |
| male_ID | Correlated random slope | stimulus_call_OP_ordinal_facIntermediateOP | 39 | 0.17 | 0.41 |
| male_ID | Correlated random slope | stimulus_call_OP_ordinal_facHighOP | 39 | 0.27 | 0.53 |
| male_ID | Correlated random slope | Motif_preceding_stim_call300ms_tone | 39 | 3.32 | 1.82 |
| male_ID | Correlated random slope | Motif_preceding_stim_call600ms_ol_only | 39 | 5.4 | 2.32 |
| male_ID | Correlated random slope | Motif_preceding_stim_call600ms_ol_synch | 39 | 7.51 | 2.74 |
| male_ID | Correlated random slope | Motif_preceding_stim_call600ms_synch_only | 39 | 5.64 | 2.38 |
| male_ID | Correlated random slope | Motif_preceding_stim_call600ms_tone | 39 | 4.83 | 2.2 |
| male_ID | Uncorrelated random slope | Trigger_call_elaboration_score | 39 | 0.05 | 0.23 |
| male_ID | Uncorrelated random slope | Stimulus_delay | 39 | 0.12 | 0.35 |

Response_Delay_LMM

| Group | RE Type | Associated Fixed Effect | # Levels | Variance | Std. Dev. |
|---|---|---|---|---|---|
| male_ID | Intercept | - | 39 | 0.03 | 0.17 |
| male_ID | Correlated random slope | Response_call_onset_time | 39 | 0.001 | 0.04 |
| male_ID | Correlated random slope | stimulus_call_OP_ordinal_facIntermediateOP | 39 | 0.006 | 0.08 |
| male_ID | Correlated random slope | stimulus_call_OP_ordinal_facHighOP | 39 | 0.006 | 0.07 |
| male_ID | Correlated random slope | Motif_preceding_stim_call300ms_tone | 39 | 0.004 | 0.07 |
| male_ID | Correlated random slope | Motif_preceding_stim_call600ms_ol_only | 39 | 0.004 | 0.06 |
| male_ID | Correlated random slope | Motif_preceding_stim_call600ms_ol_synch | 39 | 0.009 | 0.09 |
| male_ID | Correlated random slope | Motif_preceding_stim_call600ms_synch_only | 39 | 0.004 | 0.06 |
| male_ID | Correlated random slope | Motif_preceding_stim_call600ms_tone | 39 | 0.02 | 0.14 |

Within_Male_Call_Elaboration_LMM

| Group | RE Type | Associated Fixed Effect | # Levels | Variance | Std. Dev. |
|---|---|---|---|---|---|
| male_ID | Intercept | - | 39 | 0.001 | 0.1 |
| male_ID | Correlated random slope | Response_call_onset_time | 39 | 0.04 | 0.19 |
| male_ID | Correlated random slope | stimulus_call_OP_ordinal_facIntermediateOP | 39 | 0.004 | 0.07 |
| male_ID | Correlated random slope | stimulus_call_OP_ordinal_facHighOP | 39 | 0.004 | 0.07 |

Between_Male_Call_Elaboration_LMM

| Group | RE Type | Associated Fixed Effect | # Levels | Variance | Std. Dev. |
|---|---|---|---|---|---|
| male_ID | Intercept | - | 39 | 0.09 | 0.3 |
| male_ID | Correlated random slope | Response_call_onset_time | 39 | 0.014 | 0.12 |
| male_ID | Correlated random slope | stimulus_call_OP_ordinal_facIntermediateOP | 39 | 0.00005 | 0.007 |
| male_ID | Correlated random slope | stimulus_call_OP_ordinal_facHighOP | 39 | 0.0004 | 0.02 |
| male_ID | Uncorrelated random slope | Stimulus_delay | 39 | 0.035 | 0.18 |

**Table SI5**: Random effects estimates from our final models. Male ID was initially nested within night ID, but testing night explained almost no variation and caused convergences issues, so it was removed. We initially included correlated random slopes for all fixed effects (except mass and water temperature for which each male only experienced a single value). However, when including correlations between random slopes and intercepts resulted in convergence issues and singular fits, we removed correlation terms, and then removed random slopes entirely if their inclusion persisted in producing such warnings.

SI6: Restricted Overlap GLMM Results

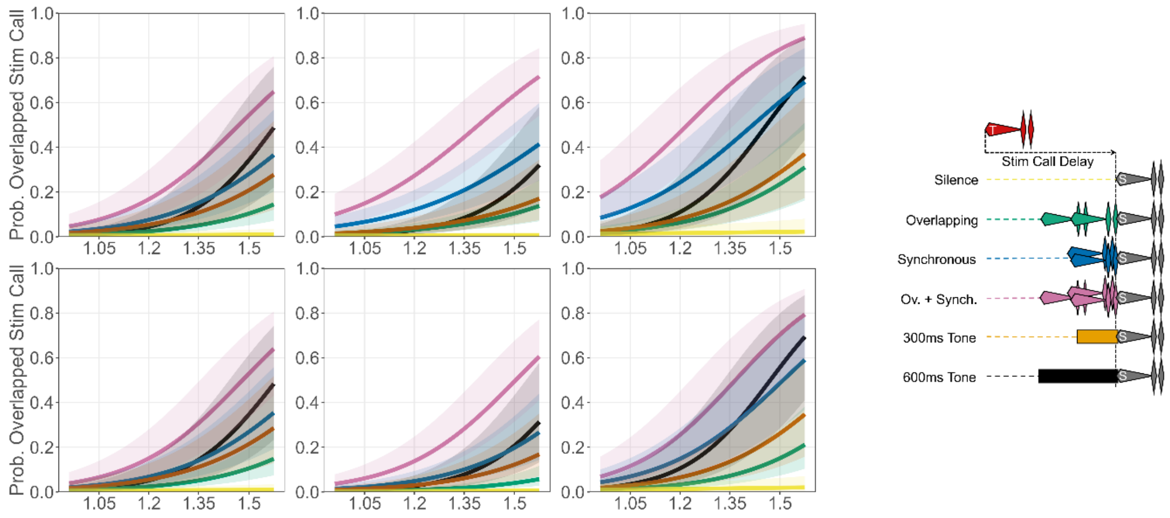


Figure SI6: Comparison of key results of *Overlap_GLMM* (Top), and *Restricted_Overlap_GLMM* (bottom). See text for details.

SI7: Methodology for calculating Inhibition Trajectories for Stimulus Calls

In figure 5, in text, we show fundamental frequency trajectories, RMS amplitude trajectories, and relative inhibition trajectories throughout the whines of our different stimulus calls. Here we explain how these relative inhibition trajectories were obtained. The code for doing so is also available in the provided Figures_Main_Text.rmd file.

In a previous paper (Larter & Ryan, 2024, Proc. B.), we tested how much males were inhibited from producing calls by tones of different frequency and amplitude combinations. This then gave us information on how inhibitory many different frequency and amplitude combinations were. As such, if we extract the fundamental frequency from a whine, as well as its RMS amplitude profile using the same window size and step length while processing, we can decompose this whine into a series of amplitude and frequency combinations unfolding through time. We can then use the model from this previous paper to predict how inhibitory these different combinations are, i.e. project these combinations into the model of male sensory tuning generated in that paper. From this, we can construct an inhibition trajectory throughout a whine. This is illustrated in Figure 2 in (Larter & Ryan, 2025).

The precise way we asked this question in this previous paper was; what is the probability that tone A is called in when embedded in tone B, with high probabilities that tone A is called in when embedded in tone B indicating that the frequency and amplitude combination of tone A made it much less inhibitory than tone B. The terminology we used was that, in this example, tone B would be the ‘reference tone’ in which tone A (the ‘test tone’) was embedded. Thus, our model compared the relative levels of inhibition between two tones; the reference tone and the test tone, with high probabilities that the test tone is called in indicating that the test tone is much less inhibitory than the reference tone. As such, we could only get predictions from our model of how inhibitory a given frequency and amplitude combination was *relative to a reference tone*. As tones of 82dB (our highest amplitude) and 1000Hz tones were the most inhibitory overall in that paper, we just set this as the reference tone for all predictions, and set the different frequency and amplitude combinations extracted from along the course of our whine as our test tones. From our model we then obtained predicted probabilities that each hypothetical test tone representing each frequency/amplitude combination throughout our whine would be called in relative to this highly inhibitory reference tone. Thus, the most inhibitory combinations from our whine would have the lowest probabilities of being called in relative to this reference tone, and the least inhibitory would have the highest probabilities. To make higher values correspond to more inhibitory combinations, we subtracted these probabilities from 1 to get an inhibition score, with higher values indicating more inhibitory combinations. As these units were somewhat arbitrary, we also normalized these scores such that the highest score within each stimulus ID whine was 1. This then gave us the relative inhibition trajectory throughout whines that arises due to the concurrent changes in frequency and amplitude.

SI8: Interesting Patterns in Latencies of Overlapping Calls

In Figure 5, in-text, it can be seen that for intermediate OP and high OP stimulus calls there are 2 peaks evident in the histograms of overlapping response call onsets occurring in the latter parts of the whines; just before the first chuck, and one peak slightly before this (at around 0.25s). We did not expect to see these multiple peaks, so we looked more closely at what might be causing them. These seemed to mostly arise because there is inter-male variation in when overlapping calls tend to occur (see figure SI7, below), with some males tending to call during the first peak, and some during the second. This seems unrelated to body size (figure SI8).

The reasons for this bimodal response are unclear. It could be that males exhibit bimodal response latencies to the same call trigger occurring shortly after whine onsets. However, this seems unlikely due to the continuous and varied nature of response delays. A more likely explanation is that males may be responding to different call triggers present throughout the quite noisy amplitude and inhibition profiles of these particular calls (figure 5, in-text; figure SI8). If this is the case, then the two peaks may arise due to some males responding to one call trigger and some to another, but with a similar distribution of latencies. Precisely identifying the call triggers here is impossible due to the complex patterns in stimulus call whines and variation in response delays within and between males. However, we can rule out one potential call trigger; the offset of the tones or motifs preceding the stimulus call. In a previous study (Larter & Ryan, 2024), the modal response latency to the offsets of 82dB 1000Hz tones (identical to the tones used to construct tonal motifs here) was 67-69ms, with the latency peak ending at ~120ms. This is much shorter than the response latencies of overlapping calls relative to preceding tone motif offsets here, which are almost all >200ms into stimulus calls. Thus, it does seem that the most likely explanation is that different parts of the noisy whine trajectories of these calls may be the most common call triggers for different males, though some combination of call trigger identity and typical response latencies could be at play.

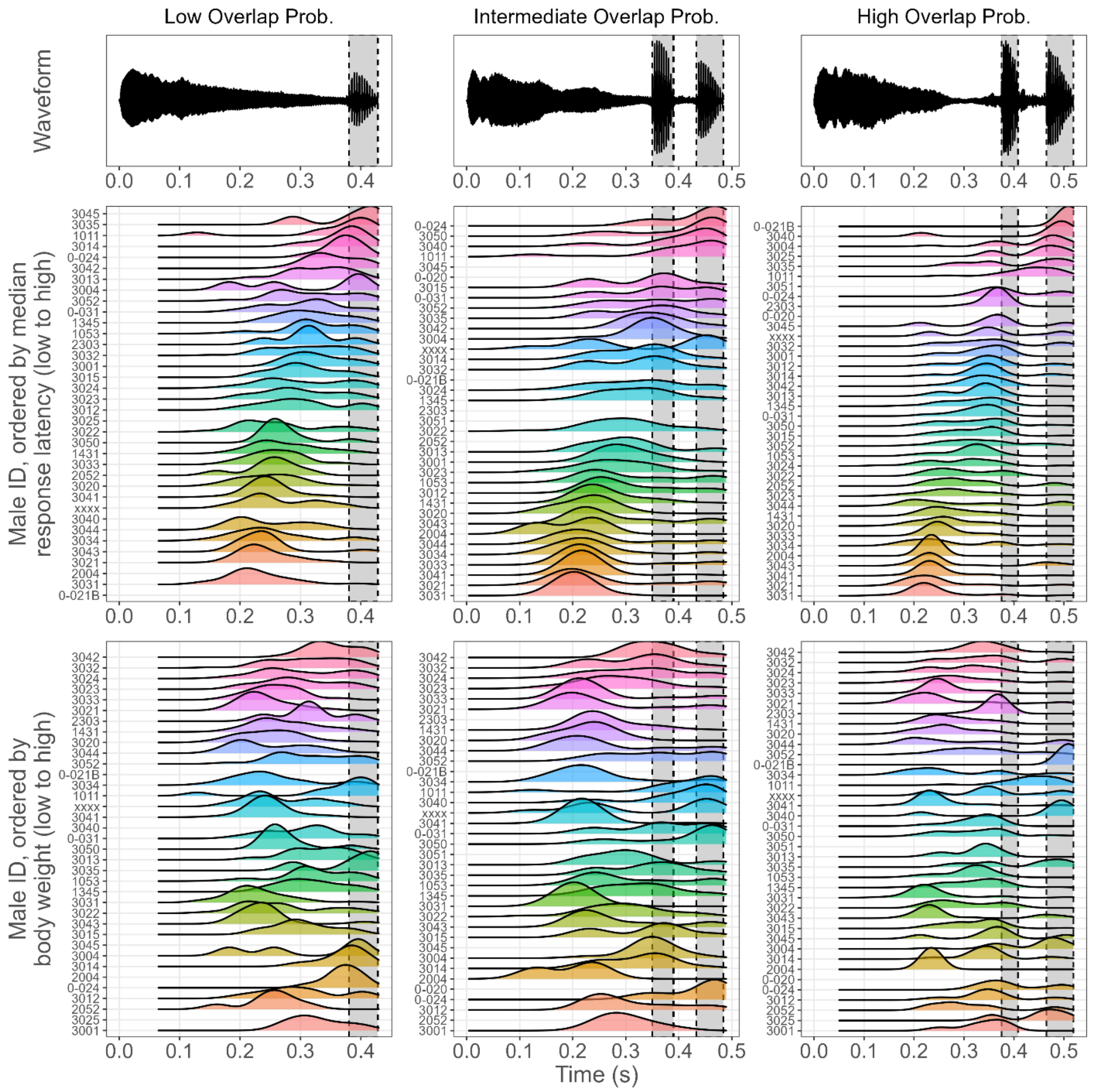


**Figure SI8**: Density plots for when onsets of overlapping calls occurred during different stimulus calls, pooled across all preceding motifs and shown separately by male ID. Blank rows indicate males produced too few overlapping calls to generate densities. In the upper density plot, males are ordered by median response latency. In the lower plot, males are ordered by body weight; as can be seen, there are no clear patterns relating to body weight. Thus, males seem to exhibit somewhat consistent differences in when overlapping calls occur throughout the same stimulus calls. This could be due to bimodal response latencies, or similar latencies in response to different call triggers. This latter explanation seems most likely (see discussion above), though some combination of both could be at play.

SI9: All Response Delays to Stimulus Calls

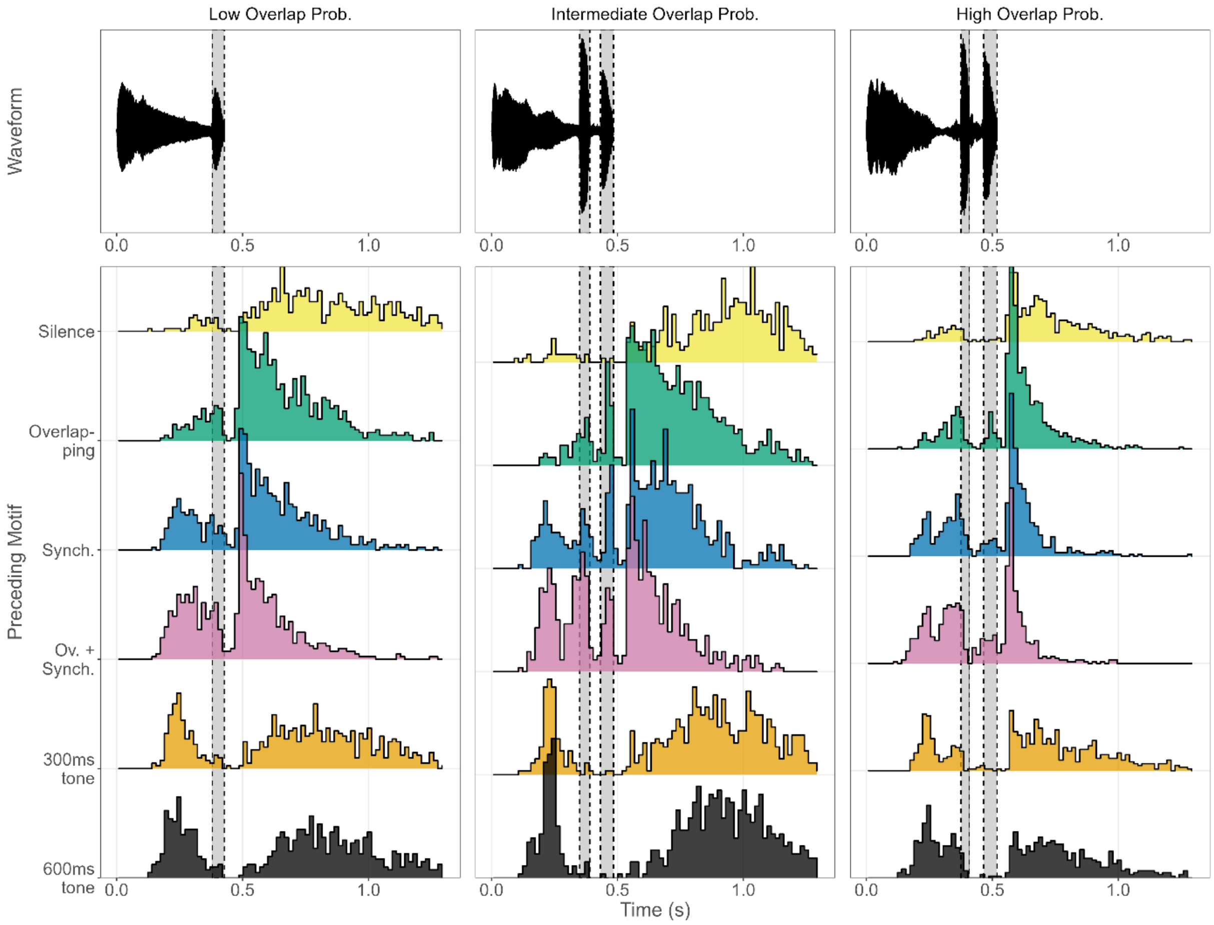


Fig SI9: Histograms of all response delays to stimulus calls (20ms bins), by preceding motif. A sharp spike shortly (~60ms) after stimulus call offsets is visible when stimulus calls are preceded by conspecific motifs, suggesting that inter-call conspecific stimulation alters call-timing mechanisms such that call offsets trigger very short latency calls (near the lower limits of this species' response times, which seems to be ~50ms: Larter & Ryan, 2024). No such tight spiking pattern is visible for responses to calls preceded by silence or tones.

SI10: Call Elaboration Distributions for all Males

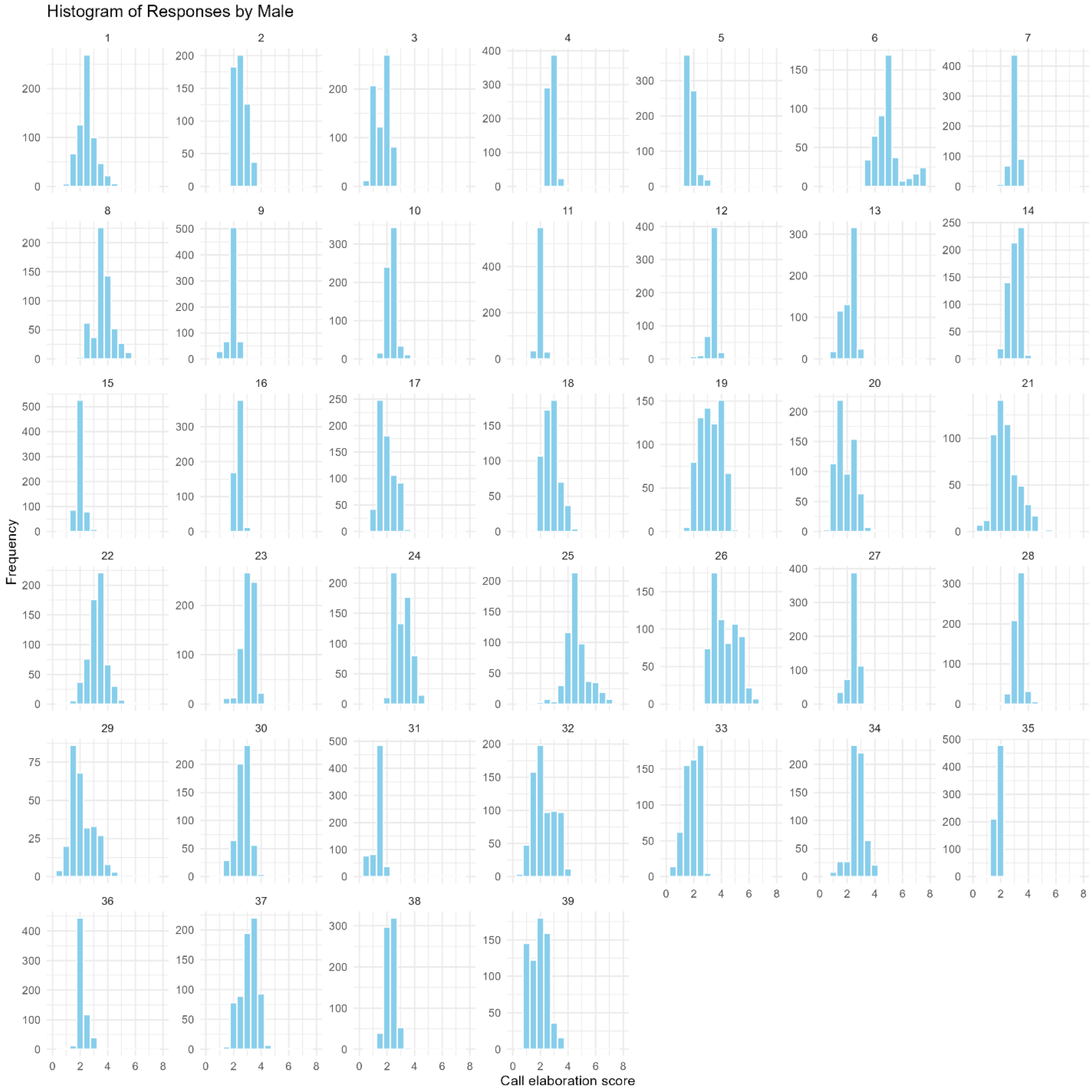


Fig SI10: Distributions of call elaboration scores throughout trials for all male subjects used.
